# Ecological specificity and interconnectivity in Danish environmental resistomes

**DOI:** 10.64898/2026.07.09.737170

**Authors:** Yu Yang, Connor Lee Brown, Lei Liu, Mantas Sereika, Thomas Bygh Nymann Jensen, Mads Albertsen, Per Halkjær Nielsen, Caitlin Margaret Singleton

## Abstract

The environment plays a critical role in “One Health”, facilitating the evolution, persistence, and dissemination of antibiotic resistance genes (ARGs). However, environmental resistomes have primarily been characterized through scattered studies of individual habitats, limiting our understanding of baseline resistome structure and connectivity across landscapes at a national level. Here, we analyzed metagenomes from 7,000 Danish environmental samples (24 Tbp), collected from 21 distinct habitats including soils, sediments, aquatic environments and wastewater treatment plants (WWTPs). We identified core ARGs for establishing the environmental baseline, and habitat-associated indicator ARGs for facilitating source tracking. Using an additional 110 deep long-read metagenomes (9 Tbp data), we found a subset of cross-habitat commonly abundant ARGs was carried by diverse hosts and associated with mobile genetic elements, suggesting their potential role in resistome connectivity across ecosystems. Additionally, although natural habitats had much lower resistome relative abundance and transferability than human-associated habitats, some mobile environmental ARGs exhibited links to those in human pathogens. Our results support the implementation of environmental resistome surveillance by providing an environmental ARG baseline, enabling future ARG source-tracking and cross-habitat ARG dissemination assessment.

## Introduction

Antibiotic resistance (AR) has been recognized as a growing threat to public health for many years ^1^. In Europe, annual deaths attributable to AR increased 26% over three years from 30,730 in 2016 to 38,710 in 2019 ^2^. Globally, 1.27 million deaths were directly attributable to bacterial AR in 2019 ^3^. Without action, the death toll is anticipated to overtake cancer ^4^. AR is ancient ^5^ and many clinically-relevant antibiotics are natural products synthesized as agents in presumed microbial chemical warfare. For example, glycopeptide biosynthesis and the associated self-resistance evolved approximately 150-400 million years ago in Actinobacteria ^6^. While AR originates from microbial ecology and evolution ^5^, the current AR crisis, including its global spread and clinical impact, has been exacerbated by societal and environmental systems.

One key driver of the spread of AR is horizontal gene transfer (HGT) of antibiotic resistance genes (ARGs), determinants of resistance in bacteria. The ability of pathogens to rapidly acquire ARGs via HGT ^7^ has resulted in a growing demand for the development of new antibiotics, a process that is beset with practical and economic challenges ^8^. Another major force underlying the spread of AR is the interconnection of human, animal, and environmental sectors, as summarised by the “One Health” framework. In brief, AR in one arena can be transmitted and propagated into another because of their inherent interconnectivity. For example, ARGs found in non-human settings can be acquired by human pathogens (for example, via human consumption of food animals), resulting in elevated rates of resistance ^9,10^. To combat these risks, it is increasingly apparent that a concerted, systems-level effort that considers all sectors is required ^11^.

While substantial effort has been made to characterize AR in humans and animals in Denmark ^12,13^, environmental AR has rarely been addressed as a systemic problem. The environment is a key player in the evolution and dissemination of ARGs ^7^, and is a reservoir for ARG acquisition and potential exchange with clinical pathogens ^14,15^. Previous studies have demonstrated the presence of near-identical ARGs encoded by soil bacteria and human pathogens. For example, one study observed near-identical copies of extended-spectrum beta-lactamase (ESBL) *CTX-M* ^16^ and quinolone resistant *qnrA* ^17^ in clinical *Kluyvera* and environmental *Kluyvera* and *Shewanella* isolates, while another study found seven soil-derived ARGs shared 100% sequence identity to ARGs encoded by human pathogens ^14^. As such, monitoring of environmental AR is important for building up the needed knowledge base for systems-level decision making, which includes environmental impact assessment or stewardship practices. However, environmental ARGs differ markedly in abundance, host range and mobility. Therefore, key monitoring objectives are needed for effective practices: 1) assessing the baseline-prevalence and distribution of ARGs; 2) defining environmentally differential ARGs (which can be used for source tracking or to identify leakage between specific environments); and 3) identification of priority ARGs at risk of mobilization across habitats. In sum, these measures can serve as a knowledge base to guide stewardship practices and intervention efforts by identifying key nodes of control, limiting use of at-risk antibiotics, and contributing to databases of potentially novel or emerging ARGs across various environmental habitats.

Here, we report analysis of the environmental resistome of Denmark and explore its features using a large-scale and geographically-dense sampling of 21 environmental habitat classes from the Microflora Danica (MFD) dataset ^18^. Using 7,156 metagenomes, we present ARG diversity and abundance, and compare within- and between-habitat resistomes to identify a small but highly abundant fraction of core ARGs (i.e., presence in all samples in a habitat classification). In order to facilitate systems-level insights about interconnectivity, we identify indicator ARGs distinguishing environments, presenting potential targets for source-tracking. Deep long-read metagenomes for 110 samples ^19,20^ revealed the genetic context and host range of commonly abundant ARGs across habitats. Through comparative genomics using these and other public genomes, we assess the HGT potential among human-, animal-, and environmentally-derived genomes, comparing different One Health sectors. This study provides an ecological framework for understanding the baseline ARGs that persist in the environment, ARGs that are characteristic of a habitat, and ARGs that are of higher potential in moving across habitats. Together, these insights provide a foundation needed to help inform future AR management efforts and to reveal how the environment is involved in ARG dissemination.

## Methods

### Sampling, DNA sequencing and quality control, and datasets acquisition

In total, 7,156 sequenced samples were collected from environmental habitats across Denmark through collaborative sampling efforts in the Microflora Danica project (MFD) ^18^. These samples represent a broad range of natural and urban landscapes to map the national resistome profile (SI1 S1 Supplementary Methods). Detailed sampling, DNA extraction, DNA sequencing, and sequencing read quality control are described previously ^18^. Briefly, extracted DNA samples were sequenced on the Illumina NovaSeq 6000 platform (PE150) producing a median depth of 5 Gbp and quality controlled using fastp ^21^ (v0.23.2) (see SI1 Figure S1 for workflow).

As explained in the original publication ^18^, each metagenome was associated with a detailed habitat description, detailing the “Sample type” (i.e., soil, sediment, and subterranean), “Area type” (i.e., natural, urban, and subterranean), and 3-level habitat ontology providing as much habitat information as we could obtain. An example of a 3-level habitat ontology of a “Sediment” “Urban” is MFD ontology level 1 (MFDO1): Freshwater; MFDO2: Standing freshwater; MFDO3: Lake, Rich pondweed. Our analyzed samples spanned a total of 21 habitat classification at the MFDO1 level (combining Sample type and Area type descriptions), and it is the habitat level this work is based on, unless otherwise specified. Additionally, to address the effect of spatial autocorrelation on resistome, subsets of spatially thinned datasets generated by the original study ^18^ using the 1-km and 10-km reference grids of Denmark (with a total of 3,947 and 1,718 samples, respectively) were used to repeat the ordination analysis.

To gain insights from the global and One Health perspectives, published short-read metagenomes were collected for ARG abundance comparison between Danish and non-Danish samples (SI2 Table S1) including non-Danish wastewater treatment plant (WWTP) influents (n=21), activated sludges (AS) (n=20) and pig guts (n=20), as well as Danish human gut (n=20) and pig guts (n=20). These public metagenomes were processed in the same way as the MFD short-read metagenomes. In addition, we examined 110 Danish long-read metagenomes (SI2 Table S2) to investigate ARG hosts and genetic backgrounds. The metagenome assemblies and metagenome-resolved genomes (MAGs) of medium- (MQ) to high-quality (HQ) were used. These datasets included the MFD project ^19^, with 87 deep Nanopore metagenomes covering 10 habitats of both urban and natural soils and sediments, and the Danish MiDAS project ^20^, with 23 hybrid Illumina-Nanopore metagenomes of Danish AS. Furthermore, to compare HGT potential in different compartments of the One Health cycle, habitat-specific genomes were downloaded including 1,015 soil genomes from RefSoil ^22^, 3,733 Danish activated sludge genomes ^20^, 3,801 human pathogen genomes from BV-BRC ^23^ (downloaded on Jan 11 2023 by searching keyword “pathogen” with filters of “Good” genome_quality and “Human” Host_common_name), 4,744 human gut genomes ^24^, as well as pig gut (1,376) and marine (1,496) representative genomes from MGnify database (v1.0 species catalogue) ^25^. The representative genomes from GTDB ^26^ (R214) were also acquired for searching the general host range of identified ARGs.

### Characterization of ARGs and mobile genetic elements (MGEs)

For quality-trimmed Illumina short-read metagenomes, ARGs were detected and quantified using ARGs-OAP ^27^ (v3.0). As the bacterial and archaeal fraction can vary largely across samples from different environmental habitats ^18^, outputted ARG abundances in rpkm were further normalized to the microbial fraction of the sequencing data, thus reporting reads per kilobase per million mapped microbial reads (rpkm microbial reads). Microbial fraction was predicted using the “microbial_fraction” module of SingleM ^28^ (v0.16.0, GTDB r214 metapackage). Resistome profiles were visualized using heatmaps and the ampvis2 ^29^ R package (v2.8.9). The sharedness and uniqueness of detected ARGs in different habitats were also investigated using ampvis2 ^29^ (v2.8.9, function *amp_core*) and visualized using the ‘ComplexUpset’ R package ^30^ (v1.3.3).

ARGs on quality-trimmed Nanopore (ONT) long reads, assemblies, MAGs, and habitat-specific genomes were first searched against the structured antibiotic-resistance gene database (SARG, v3.0) using DIAMOND blastx ^31^ (v.2.1.6) at minimum cutoffs of 80% identity and 80% subject coverage. Predicted ARG-carrying ONT long reads were then frameshift-corrected using proovframe ^32^ (v0.9.8). Identified ARG-carrying frameshift-corrected long reads, assemblies, MAGs, and representative genomes were then searched for transposases, integrases, and recombinases associated with MGEs using DIAMOND blastx (80% identity, 60% subject coverage, following previous studies ^33,34^) against MGE databases comprising both ISfinder-sequences ^35^ (downloaded on Mar 3 2023) and a text-mined collection of UniRef100 sequences in UniProt ^36^ (downloaded on Mar 3 2023) with keywords of “Integrase OR recombinase OR transposase” in annotation. Overlapping MGE annotations were merged to remove duplicate annotations and the annotation with the lowest evalue from BLAST were kept using the R package ‘GenomicRanges’ ^37^ (v1.50.2, function *reduce*).

### Detection of core and indicator ARGs

Core and indicator ARGs are particularly beneficial for long-term ARG monitoring, source-tracking, and management. Core ARGs of a habitat were defined as those found in 100% samples of that habitat ^38,39^. Candidate indicator ARGs were identified by the ‘indicspecies’ package ^40^ (v1.8.0, *function multipatt*) using the ARG abundances in rpkm corrected by microbial fraction as input data matrix (delug=TRUE, statistical significance tested by permutation test of 999 permutations, and Benjamini-Hochberg corrected *p*-value <0.05). The association between an ARG and a biome was outputted as an indicator value, which is the product of detection specificity and sensitivity. Identified ARGs with a minimum indicator value of 0.5 (0-1, higher value means stronger indicator) and a minimum 0.5% relative abundances in all samples of a biome were characterized as indicator ARGs for that specific biome.

### Prediction of the phylogenic hosts and genetic background of ARGs

ARGs were only linked to their microbial host through MAGs or genomes. To identify the overall host range of an ARG, representative genomes in GTDB ^26^ (R214) were used. Danish MAGs were taxonomically classified using GTDB-tk (based on GTDB r214) as part of the MAG construction pipeline mmlong2-lite ^19^ (https://github.com/Serka-M/mmlong2-lite, v1.0.2). Assessment of the genetic location of ARGs (chromosomal, plasmidial, viral, and unknown) relied on ARG-carrying assemblies with a minimum 1kb length filtered by seqkit ^41^ (v2.5.1). ARG-carrying contigs were first characterized to be plasmid-born or virus-borne by geNomad ^42^ (v1.8.0; end-to-end --enable-score-calibration, with the default post-classfication filters), plasflow ^43^ (v1.1). GeNomad-predicted plasmids were further filtered to include only those cross-validated by plasflow ^43^ (v1.1). Resulting predicted plasmids were then classified by Plascad ^44^ (v.1.17) to determine plasmid mobility types (i.e., mobilizable, conjugative, non-mobilizable plasmids) through the HMM based search of plasmid transfer associated genes. GeNomad-predicted viral contigs were quality-assessed by checkv ^45^ (v1.0.1 with v1.5 db) to exclude those with “Not-determined” quality or found with no viral proteins by checkv. “Chromosomally”-located ARGs are charaterized as those found in bins, while excluding those found on predicted plasmidial and viral contigs. Estimation of the per habitat level (i.e., AS, soil, and sediment) ARG’s genetic background (i.e., viral, plasmidial, and chromosomal) for each ARG type was performed considering both the coverages of ARG-carrying contigs and total datasize of metagenomes and was given in the median of each sample’s contig-coverage-corrected ARG hits per Gb raw metagenome data per habitat.

### Measurement of ARG HGT potential within each habitat

To investigate the ARG transfer potential in different sectors of the One Health cycle, we used MAGs and assemblies from Danish environmental samples, as well as published habitat-specific genomes of the environment and guts of healthy humans and farmed pigs. The mobility potential of ARGs was analyzed using the colocalization of ARGs and MGEs on assemblies and genomes, where the number of MGEs (i.e., transposases, integrases, and recombinases) within the assigned radii (0-10kb) ^46^ from each ARG were counted. For each identified ARG on a genome/MAG from a habitat, the distance between the midpoints of the seed ARG and each identified MGE on this genome were recorded. Distances were rounded to the nearest 100bp as a stepsize and filtered to a maximum of 10kb (for example, 10kb represents 100 steps). The counts of MGEs at different distances (in stepsize) to this seed ARG were counted sequentially after ranking the distances to the ARGs ascendingly. Then, for each habitat, taking the median counts of MGEs for each distance to ARGs in stepsizes to represent the incidence of MGE for ARG at different distances.

### Determination of shared ARGs between the environment and human pathogen

ARG sequences in the frameshift-corrected ONT reads of Danish samples and in the human pathogen genomes were extracted using seqtk (https://github.com/lh3/seqtk, v1.4, *subseq* subcommand) and BEDtools ^47^ (v2.30.0) for the identification of potentially transferable ARGs between Danish environment and human pathogens. To do so, we compiled a BLAST database of all seqtk-extracted ARGs identified in the human pathogen genomes, then performed all-versus-all BLASTn on all ARG sequences found on the frameshift-corrected ONT reads of Danish samples. Stringent cutoffs of a minimum of 98% identity and 98% subject coverage were used to determine shared ARGs between the Danish environment and human pathogens. Colocalization of ARGs and MGEs were visualized using the R package ‘gggenes’ ^48^ (v0.5.0).

### Statistical analyses

All statistical analyses were performed in R ^49^ (v4.4.0) and all plots were generated by the ‘ggplot2’ ^50^ package (v3.5.2). To perform Principal Coordinate Analysis (PCoA) for the resistome in different habitats, the relative abundances of ARG subtypes in all samples were first Hellinger transformed using the ‘vegan’ package ^51^ (v2.6-10 function *deconstand*), then Bray-Curtis distances were calculated based on the transformed abundances by the ‘parallelDist’ package ^52^ (v0.2.6, function *parDist*), and lastly, the resistome profiles for samples were visualized in PCoA using the ‘ape’ package ^53^ (v5.8, function *pcoa*). Permutational multivariate analysis of variance (PERMANOVA) was applied to test resistome dissimilarity among sample types or habitats (*p* < 0.05) using the ‘vegan’ package ^51^ (v2.6-10, function *adonis2*) with 999 permutations. Microbial profiles at the genus level were generated from the original study ^18^, which were based on the 16S rRNA gene fragments extracted from the metagenomic reads. Ordination analysis for the microbial community was performed the same way as the resistome profiles. To evaluate the concordance between microbial communities and resistome profiles, Procrustes analysis (function *procrustes* and *protest*, 9999 permutations) and Mantel tests (based on Spearman’s rank correlation, 999 permutations, function *mantel*) were performed using the calculated Bray-Curtis distance matrices with the ‘vegan’ package ^51^ (v2.6-10). Kruskal-Wallis and the Dunn *post-hoc* test was conducted to compare the median of ARG abundances across different habitats using the ‘rstatix’ package ^54^ (v0.7.2, function *kruskal.test* and *dunn_test*). In the case of adding comparison *p*-values to a ggplot, the ‘ggpubr’ package ^55^ (v0.6.0) was used, for example for the spearman correlation between short-read metagenomes subsampled at different depths and Wilcoxon test for the comparison of 2 groups (significant difference at *p* < 0.05). The heatmap of indicator ARGs was plotted with ‘ComplexHeatmap’ ^56^ (v2.22.0) and clustered based on “spearman” correlation distance using the “centroid” clustering method. To determine whether human pathogens had significantly (*p* < 0.05) greater HGT potential than other habitats regardless of the increasing genetic distance from an ARG, Fisher’s exact test was done using the base R stats package ^49^ (function *fisher.test*). To evaluate whether the available sample sizes of the published short-read metagenomes collected for One Health comparisons were sufficient to resolve differences among dataset sources, an analysis of similarities (ANOSIM) based on Bray–Curtis dissimilarities of ARG abundance profiles was performed using the *anosim* function in the ‘vegan’ package ^51^ with 999 permutations.

## Results and discussion

### Danish environmental resistomes vary in abundance and diversity across habitats

As the first step towards a Danish national environmental resistome survey, we investigated ARG abundance and diversity across 7,156 shallow metagenomes from the Microflora Danica project ^18^ broadly covering 3 different sample types (i.e., soil, sediment, and water), spanning 21 habitats from urban, natural, and subterranean areas (metadata in SI3). The ARG abundances varied across habitats and area types (all Kruskal-Wallis test *p* < 0.05, Figure 1a, SI1 Figure S4). We observed one order of magnitude difference in the total ARG abundances across different habitats (Figure 1a, SI2 Table S3), with the lowest abundance in natural saltwater sediments (median 45.3 rpkm microbial reads), and the highest in wastewater influents (median 1137.9 rpkm microbial reads). The *post-hoc* analysis indicated that most sediment habitats had significantly lower ARG abundances compared to soils and waters (Dunn’s test with BH-*p.adj* < 0.05, SI1 Figure S5), which agreed with recent work of a smaller scale ^57^. Salinity was associated with ARG abundance variation, particularly in sediment and water habitats, likely through its influence on microbial community composition ^58^ and its role as an ecological barrier for microbial and resistome dispersal across habitats ^59^, thereby contributing to the observed differences in ARG abundances. (Figure 1a).

**Figure 1.**
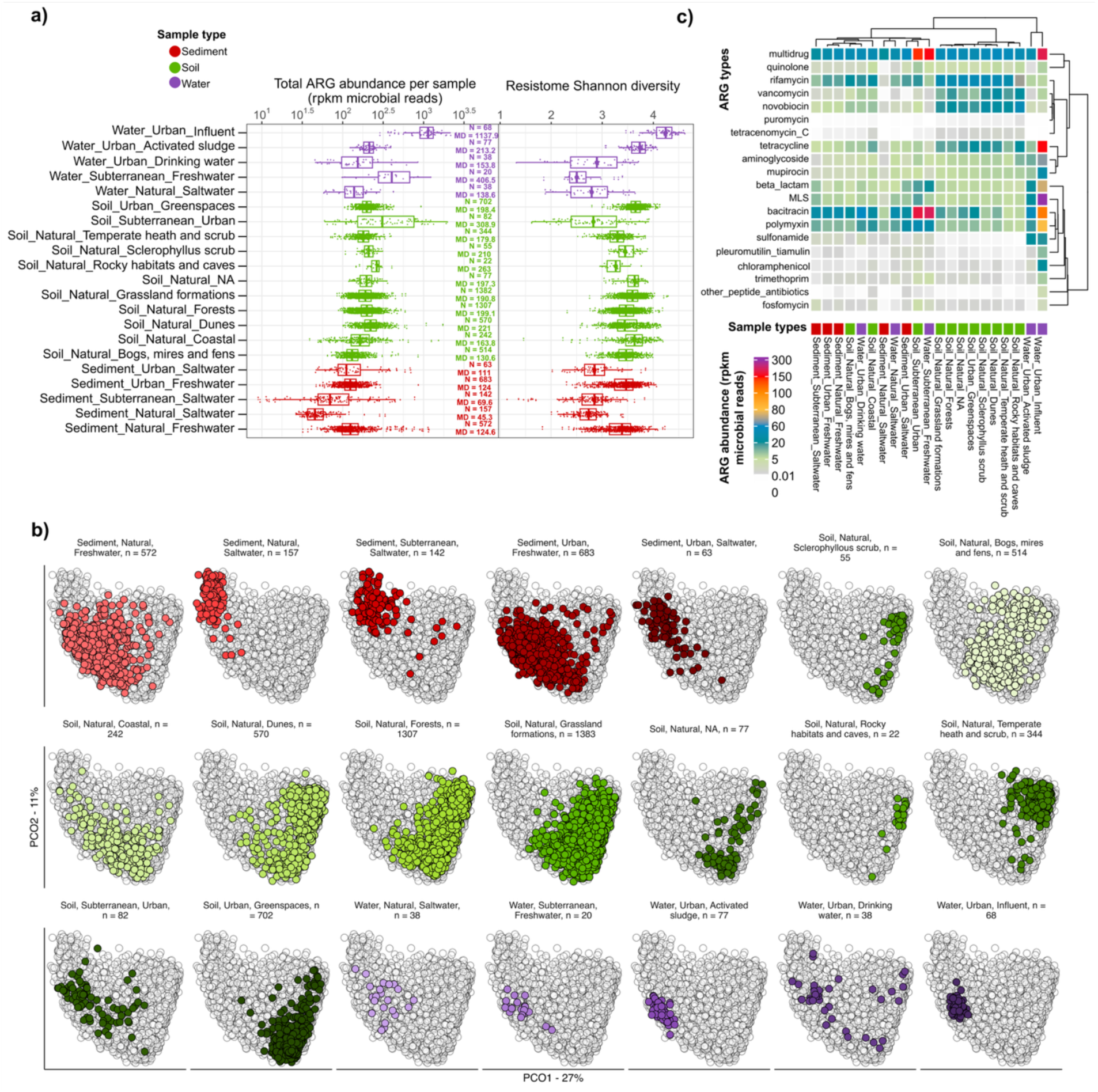
Abundance and diversity of Danish environmental resistome. **a)** Left panel boxplot showing the distribution of total ARG abundances per sample across different habitats of different sample types (Kruskal-Wallis *p* < 2.2e-16). N: number of samples, MD: median. Right panel boxplot showing the distribution of Shannon diversity of each sample across different habitats. Influent refers to WWTP influent samples. Rarefaction applied to the minimum depth of ARG-like reads; **b)** PCoA analysis of the resistomes across different environmental habitats based on Bray-Curtis dissimilarity on Hellinger-transformed relative abundances of ARG subtypes, adonis2 PERMANOVA R^2^ = 0.17 and *p*-value < 0.001 in 999 permutations; **c)** Heatmap showing the resistome composition across different habitats at the ARG type level. Dendrograms were created based on the Spearman correlation-based distance matrix and centroid clustering method. ND: non-detectable. MLS: Macrolides-lincosamides-streptogramines. Each habitat had a minimum of 20 samples.

Additionally, ARG abundances were associated with urbanization level (based on the habitat classification at the “Area type” level), as urban samples had significantly higher ARG abundances than natural samples (SI1 Figure S4b). Compared to non-Danish datasets, Danish ARG abundances in the WWTPs and farmed pig guts (SI1 Figure S6, ANOSIM, R = 0.809, P = 0.001), and human guts ^60^ were lower than those observed in other regions (two-sided Wilcoxon rank-sum test, *p*-value <0.05). Compared to other One Health sectors in Denmark, data sourced from farmed pig guts had higher ARG abundances than data sourced from healthy human guts and our environmental data (SI1 Figure S6, ANOSIM, R = 0.913, P = 0.001). The strong separation among dataset sources observed through ANOSIM indicates that between-source variation substantially exceeded within-source variation despite differences in sample numbers among habitats. However, these cross-study comparisons involved differing technical variables (sampling, sample processing, and DNA extraction methods), which could introduce variation that impacts biological inferences.

### Resistome diversity

Danish environmental resistomes generally separated based on habitat in ordination analysis (Figure 1b), and such separation explained 17% of the resistome dissimilarity (*adonis2* PERMANOVA R^2^ = 0.17, *p*=0.001, 999 permutations). Broadly, resistomes showed habitat-associated clustering among soils, sediments, and waters (Figure 1b), with partial overlap among some habitats and clear separation among others. Similar habitat-associated clustering was observed after spatial thinning (SI1 Table S7), indicating that the pattern was robust to local sampling density. In general, soil resistomes shared ordination space, with the exception of “Temperate heath and scrub” and “Bogs, mires and fens” soils. These broad habitat classifications encompass substantial variation in environmental conditions (e.g. nutrient availability, pH and moisture), which is likely to support highly heterogeneous microbial communities and, consequently, more diverse resistome profiles ^20^. This variation likely reflects underlying differences in microbial community structure, which was strongly associated with resistome composition (spatially-thinned subset in 10 km reference grid: Procrustes correlation = 0.87, protest *P* = 0.0001; Mantel *r* = 0,855, *P* < 0.001, SI1 Table S8), consistent with previous observations that microbial community composition is a major determinant of resistome composition ^61^. In addition, salinity appeared to separate resistome profiles for sediment, soil, and water samples (Figure 1b), likely reflecting salinity-driven differences in microbial community composition. Saline and non-saline sediments and waters formed distinct clusters, and coastal soils were more dissimilar to natural and urban soils. Impact of salinity on resistome abundance and diversity has also been reported in oceans ^62^. Interestingly, urban soil (i.e., “Soil, Urban, Greenspaces” with n=702) formed tighter resistome clusters compared to natural soils of comparable sample sizes (e.g., “Soil, Natural, Coastal” with n=242, “Soil, Natural, Dunes” with n=570, “Soil, Natural, Bogs, mires and fens” with n=514, Figure 1b). This pattern may partly reflect reduced microbiome diversity in human-disturbed soils compared with natural soils ^18^, potentially contributing to resistome homogenisation among disturbed soils. Together with previous studies ^63^, our findings highlight the importance of microbial diversity in limiting the spread of ARGs.

Habitat-separations based on the resistome varied among different drug types (SI4). Separations at the sample type level (i.e., soil, sediment, water) were mostly observed for beta-lactam, polymyxin, and multidrug-associated ARGs (Figure 1c, SI4). Additionally, WWTP samples not only harbored the highest resistome diversity (Figure 1a), but were also distinct from other habitats in when considering ARGs associated with commonly-used antibiotics in Denmark ^64^, including macrolide-lincosamide-streptogramin (MLS), beta-lactams, aminoglycosides, tetracyclines, sulfonamides (SI4). The broad soil categories showed substantial resistome variability across many drug classes, as reflected by their dispersion in ordination space (SI4). This variability likely reflects, at least in part, the wider environmental breadth captured by soils, including heterogeneous physicochemical conditions and microbial communities ^18^. Despite this variability, soil samples exhibited similar profiles of tetracycline, rifamycin, and multidrug ARGs, as they formed tight clusters in these drug classes (SI4). These ARGs are frequently reported in soils ^61^, which could be due to the versatility of multidrug-ARG-associated efflux pumps that can perform functions not limited to AR ^65^ and the historical and indirect tetracycline inputs into soils through livestock manure, sewage sludge land amendments ^66^, respectively. MLS ARGs, on the other hand, exhibited moderate separation power between urban and natural sources of soils and sediments, such as the separate ordination clusters formed between “Soil, Urban, Greenspaces” and other soils of natural source (SI4). This pattern indicates the potential utility of MLS ARGs as indicators of anthropogenic influence on environmental resistomes. Overall, the Danish environmental resistome is diverse and shows differentiation among ARGs targeting different drug classes.

Quantifying ARG diversity and abundances is essential for microbial AR stewardship. However, ARG identification and quantification from sequencing data alone is challenging, as some ARGs may possess functions besides resistance ^67,68^, confer AR when overexpressed ^67^, or do not directly confer resistances (e.g. regulatory genes ^68^). Although metagenomic datasets cannot directly infer phenotypic observation ^69^, they provide a standardized framework for comparing genotypic resistance potential across large numbers of environmental samples. Thus, our large-scale dataset offers insight into the diversity, abundance and habitat distribution of genes that may directly or indirectly contribute to microbial AR.

### Core ARGs for the environmental baseline resistome

Baseline assessment of ARGs in the environment can establish a factual foundation for multiple environmental monitoring objectives, including impact assessment and stewardship practices, but this baseline is largely lacking ^70^. Here, we defined the baseline resistome of Denmark using “core” ARGs, i.e., those ARGs which are consistently detected across all samples within a habitat. Core ARGs thus represent a minimal, characteristic, and representative set of ARGs with stable associations to specific habitats.

Based on short-read mapping results, we found that ARG subtypes of higher prevalence (detected in ≥80% samples in a given habitat) were generally of higher abundance compared to rarer ARGs with RPKM microbial read values typically in the range of 1-10 and 0.1-1, respectively (Figure 2a). A total of 69 core ARG subtypes (representing 0.4-11.8% of all ARGs detected in a given environment) were identified (Figure 2bce), representing the majority of total ARG abundances (7.5-81.6%, Figure 2f). These subtypes include *macB*, *mexF*, *mexW*, *mexB*, *muxB*, and *novA*, which have also been reported in other studies as core ARGs present in a variety of habitats ^71–73^. The majority of other detected ARGs (i.e., 37-83% detected ARGs, Figure 2ae) were “rare” (i.e., detected in ≤ 10% samples in a given habitat), contributing to a small fraction of the total ARG abundances (i.e., 1-7%, Figure 2f). The consistent observation of a high abundance but low diversity of core resistomes in human, engineered, aquatic, and terrestrial samples ^38,39,74^, could arise from conserved microbial physiological functions. Most of the core ARGs (44/69) encoded efflux pumps, which are intrinsic characteristics of microbial communities (e.g. in defense systems ^74^, metabolic functions (C-, S-, and P-cycling) and signaling ^15,74^ (SI2 Table S4)). These genes are ubiquitous and have a broad range of substrates ^58^, for example, *macB* has been implicated in iron scavenging ^76^. While these genes are generally considered lower risk because they are often host-restricted and involved in broader physiological functions, they can still be enriched by anthropogenic activities ^77^ and contribute to AR when overexpressed ^78^.

**Figure 2.**
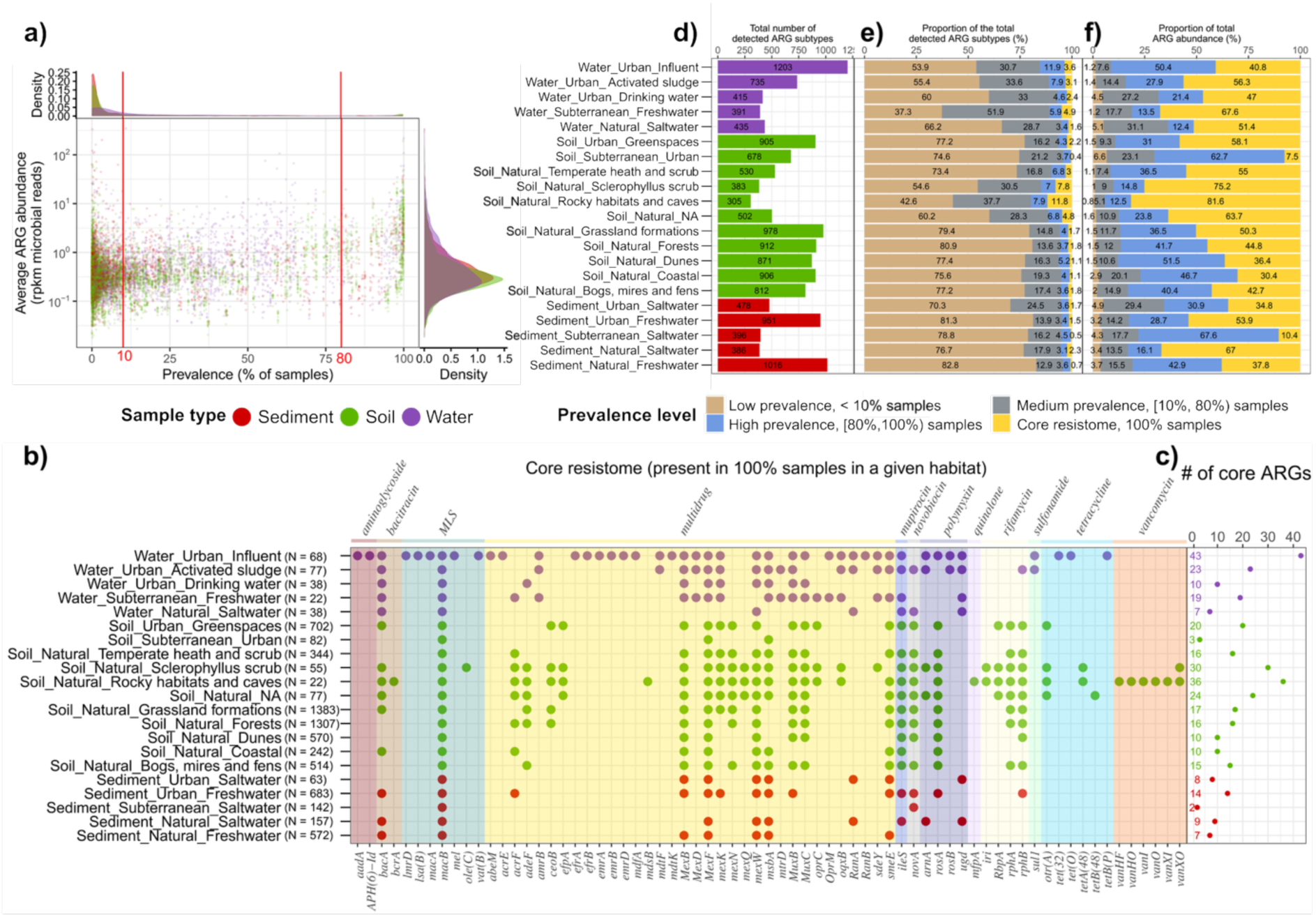
Prevalence of ARGs. **a)** Average abundance and prevalence of each detected ARG in different sample types; **b)** Detection of a total of 69 core resistomes for different habitats; **c)** The number of core ARGs in each habitat; **d)** The total number of ARG subtypes detected in each habitat; **e)** The proportion of detected ARG subtypes that were of different prevalence levels: high prevalence (present in ≥ 80% of the samples in a given habitat), low prevalence (present in ≤ 10% samples in a given habitat), and all others in between (that are present in 10-80% of samples in a given habitat); **f)** The proportion of the total ARG abundance that ARGs of different prevalence levels could account for. Influent refers to WWTP influent samples. The mean relative abundances of each ARG subtypes quantified in all samples in a given habitat were taken in calculating the contribution to the total ARG abundances by ARGs under different prevalence levels. MLS: Macrolides-lincosamides-streptogramines. Each habitat has a minimum of 20 samples.

We identified that the core resistome can reflect drug use and source microbes. Danish WWTPs had a particularly broad range of core ARGs resisting highly consumed antibiotics in Denmark ^12^, including aminoglycosides, beta-lactams, MLS, and tetracyclines (Figure 2e). The core resistome in WWTP influent overlapped with ARGs reported in the human gut ^79^, such as *macB*, *efrAB*, *msbA*, and *ileS*. This may be due to the presence of gut microbiota in the influent, which makes up 34.8% of the influent microbiome in Denmark ^80^. Compared to other sample types, soils had greater proportions of core ARGs conferring resistance to vancomycin and rifamycin (Figure 2e). This likely reflects the intrinsic ARG potential of antibiotic-producing soil core microbes, as vancomycin and rifamycin producers, e.g., *Streptomyces* and *Micromonospora*, were reported as core genera in Danish soils ^18^. Overall, the core resistome of high abundance and prevalence ARGs represents a persistent baseline of ARGs in a given habitat, with the possibility of enrichment under anthropogenic perturbations. Importantly, the core ARGs defined here likely reflect only a subset of the intrinsic resistome characteristics for each habitat. Increasing sequencing depth (or decreasing the stringency of the inclusion parameter) would likely identify more core ARGs and reveal more granular connections between environments. Nevertheless, the broad ecological coverage and extensive sampling in this study provides robustness of the core resistome characterization.

### Potential indicator ARGs suggest ecological signals

While core ARGs can define a baseline resistome, indicator ARGs offer insights into practical and cost-effective ARG targets for actionable environmental monitoring and source-tracking based on differences between habitats. The wide range of Danish environmental habitats investigated provided advantages for indicator ARG assessment at the country level. As such, we sought to identify indicator ARGs that differentiated ecological habitats. We further identified indicator ARGs that differentiate urban vs. natural sources. To facilitate source tracking, we further assigned these 21 habitats to 9 different biome clusters based on resistome clustering at the MFDO1 level (SI2 Table S5). This was possible because resistome profiles generally clustered based on habitat descriptions at the MFDO1 level, such as salinity for sediments (saltwater vs freshwater), area type for soils (soil habitat with potential petroleum pollution in “Soil, Subterranean, Urban” vs. all other soils) and stage of wastewater treatment (influent vs activated sludge) (Figure 1b). The distinction between urban and natural sources relied on the habitat ontology at the “Area type” level based on the MFD project ^18^.

Across the biome clusters, we identified 86 potential habitat indicator ARGs (with indicator value ≥ 0.5 by “indicspecies” R package ^40^, with minimum 0.5% relative abundance across all samples in a habitat) for differentiating samples from different biome clusters (Figure 3, SI2 Table S6). We focused on indicators with relative abundances above 0.5% to maximize their applicability in environmental habitat monitoring, as low-abundance ARGs are more likely to be missed by metagenomic sequencing. These signals likely reflect both habitat-associated ARGs and the distribution of their microbial hosts. Some ARGs indicating WWTP origins were mobile, and/or of medical importance, and/or found in human samples. For example, *tetO/M/W*, *OXA-129/2*, *mel*, and *ermB* are often associated with mobile genetic elements and *OXA-2* and *mel* are clinically relevant. Given the high proportion of human gut microbiota in WWTP influent (34.8% read abundances in 16S rRNA gene sequencing ^80^), it is unsurprising that some of WWTP-indicating ARGs were also found in human gut/clinical samples, such as *sul1, dfrF, tet(32),* and *tetW/O.* Enrichment of these ARGs in other habitats could suggest anthropogenic contamination. We found the bifunctional polymyxin resistance gene *arnA* to be associated with urban subterranean soils, most of which were potentially petroleum-polluted in MFD (80.5%: 66/82 samples). Interestingly, *arnA* enrichment and its association with MGEs has also been reported in petroleum-impacted marine sediments ^81^. Unlike many acquired ARGs, *arnA* has broader physiological functions in lipid A modification, contributing to cell envelope adaptation. Its enrichment therefore does not necessarily imply selection by polymyxins, but may instead reflect habitat-associated microbial communities and environmental conditions favouring organisms carrying this gene. This example illustrates that indicator ARGs should not always be interpreted as evidence of antibiotic selection, but can also serve as markers of habitat-associated microbial and resistome structure. Similarly, PNGM-1 was identified as an indicator ARG for saltwater-origin samples, consistent with previous reports of high PNGM-1 abundance in deep-sea sediments predating the antibiotic era ^82^. ARGs indicating soil origins also tended to include genes intrinsically found or chromosomally located in soil-dwelling bacteria, such as *tetV* in *Mycobacteria* and *otrA* in oxytetracyline-producing *Streptomyces*. In addition, to assess potential leakage of “urban” ARGs into “natural” environments, we identified 73 ARG indicators which differentiated the two sets (indicator value ≥ 0.5, SI2 Table S7). Of these, 17/73 ARGs were suggestive of urban origin. Within the 17 urban ARGs, eight were also indicative of specific ecological biomes. For example, a third of the urban-indicating ARGs (5/17: *lsa(B), efrA, ileS, tet(M), tetB(48)*) were indicative of WWTP influent sources. These patterns suggest that a subset of urban-associated ARGs may capture links between urban resistomes and human-associated waste streams. Together, these examples suggest that indicator ARGs often reflect habitat-associated microbial lineages and ancient or intrinsic resistome components, rather than recent antibiotic selection alone. In addition to source-tracking and monitoring, these ARGs may serve as candidate markers of cross-habitat resistome connectivity assessment, especially when interpreted together with host and MGE contexts.

**Figure 3.**
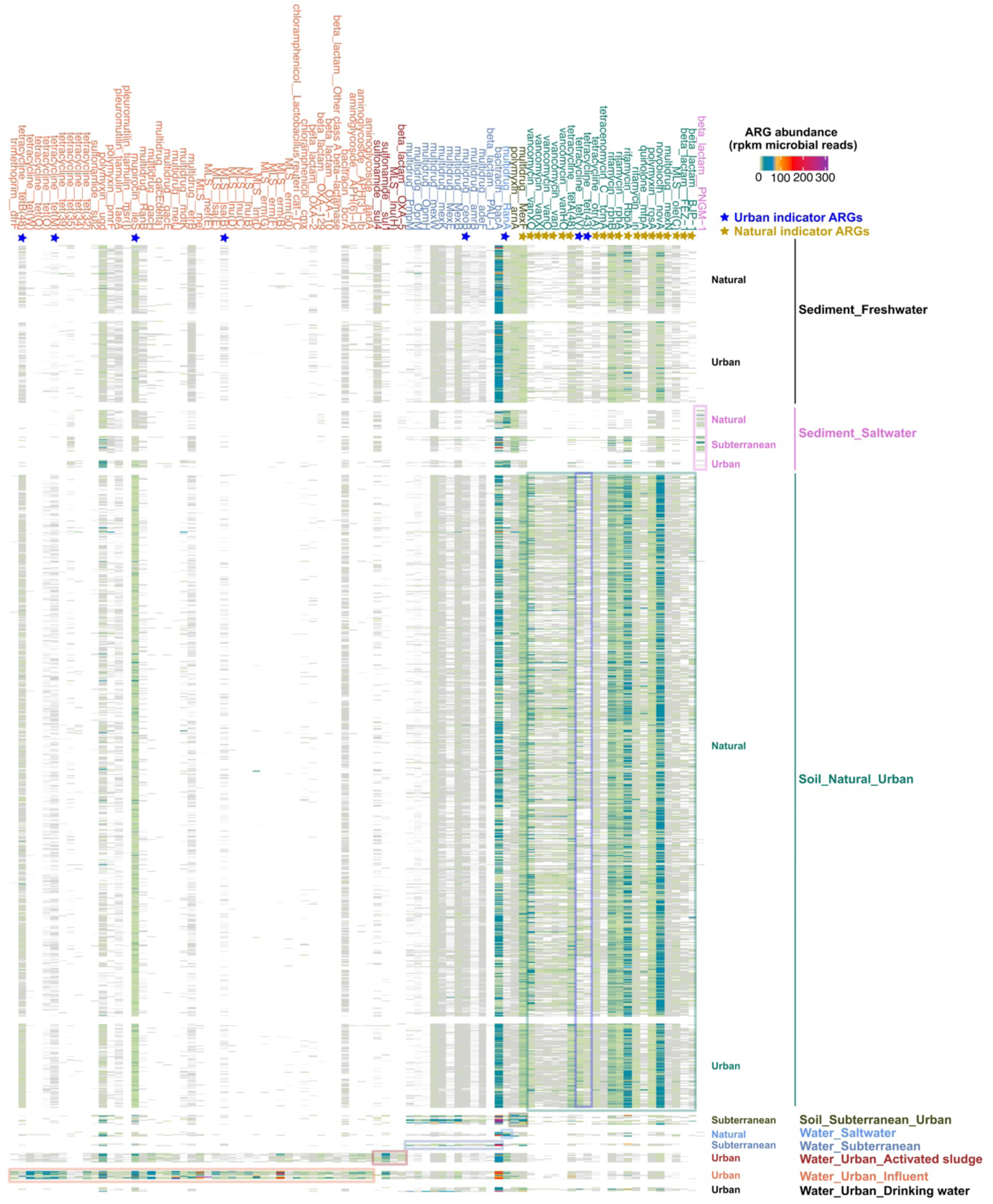
Potential indicator ARGs. Rastered heatmap showing the abundances of the 86 abundant indicator ARGs across all 7,156 samples (minimum 0.5% relative abundance in every habitat). Indicator ARGs were grouped and colored by their indicating biomes. Boxes are used to highlight the abundances of indicator ARGs in their indicating biomes. Note that both Sediment_Freshwater and Water_Urban_Drinking water biomes had no indicator ARGs, hence they were colored in black. Indicators for urban and natural habitats were also labelled as blue and yellow stars, respectively. MLS: Macrolides-lincosamides-streptogramines.

### Some ARGs show ecological connectivity

While core and indicator ARGs offer insight into the resistome similarity within habitats and differences between habitats, they do not capture the potential spread of ARGs across environments. To investigate potential ARG connectivity across habitats, we examined ARGs that are commonly abundant across habitats for their nearby MGEs and microbial hosts.

Among all the 1,810 detected ARG subtypes based on short-read metagenomes, a total of 150 ARG subtypes contributed to the top 80% of the total ARG abundance for any one habitat (SI1 Table S9, SI2 Table S8). Among these 150 ARGs, 83 were commonly abundant ARGs (found in at least two MFDO1 levels), suggesting their potential transferability across habitats (SI2 Table S8). We explored the genomic context of these 83 ARGs using Nanopore (ONT) long-read sequencing from 110 samples representing 11 habitats (87 MFD samples ^18,19^ and 23 Danish AS samples ^20^) (Figure 4a, SI2 Table S2). Most of the 83 commonly abundant ARGs found based-on short-read data were detected in the long read data (84.3%: 70/83). Of these, 68.7% (57/83) had MGE within 5 kb radius of the ARG (SI2 Table S8, Figure 4bc), compared with 47% (216/460, SI2 Table S9) for non-commonly-abundant ARGs, indicating a greater MGE association among commonly abundant ARGs and suggesting a greater potential for horizontal dissemination. A fifth (17/83) of these ARGs were frequently found next to MGE (i.e. in at least a third of the long reads from which these ARGs were identified, SI2 Table S8, SI1 Table S10, Figure 4b), suggesting their frequent association with MGEs and high HGT likelihood. CARD searches further confirmed 14/17 MGE-associated ARGs have previously been reported to be plasmid-associated ^83^ (SI2 Table S8).

**Figure 4.**
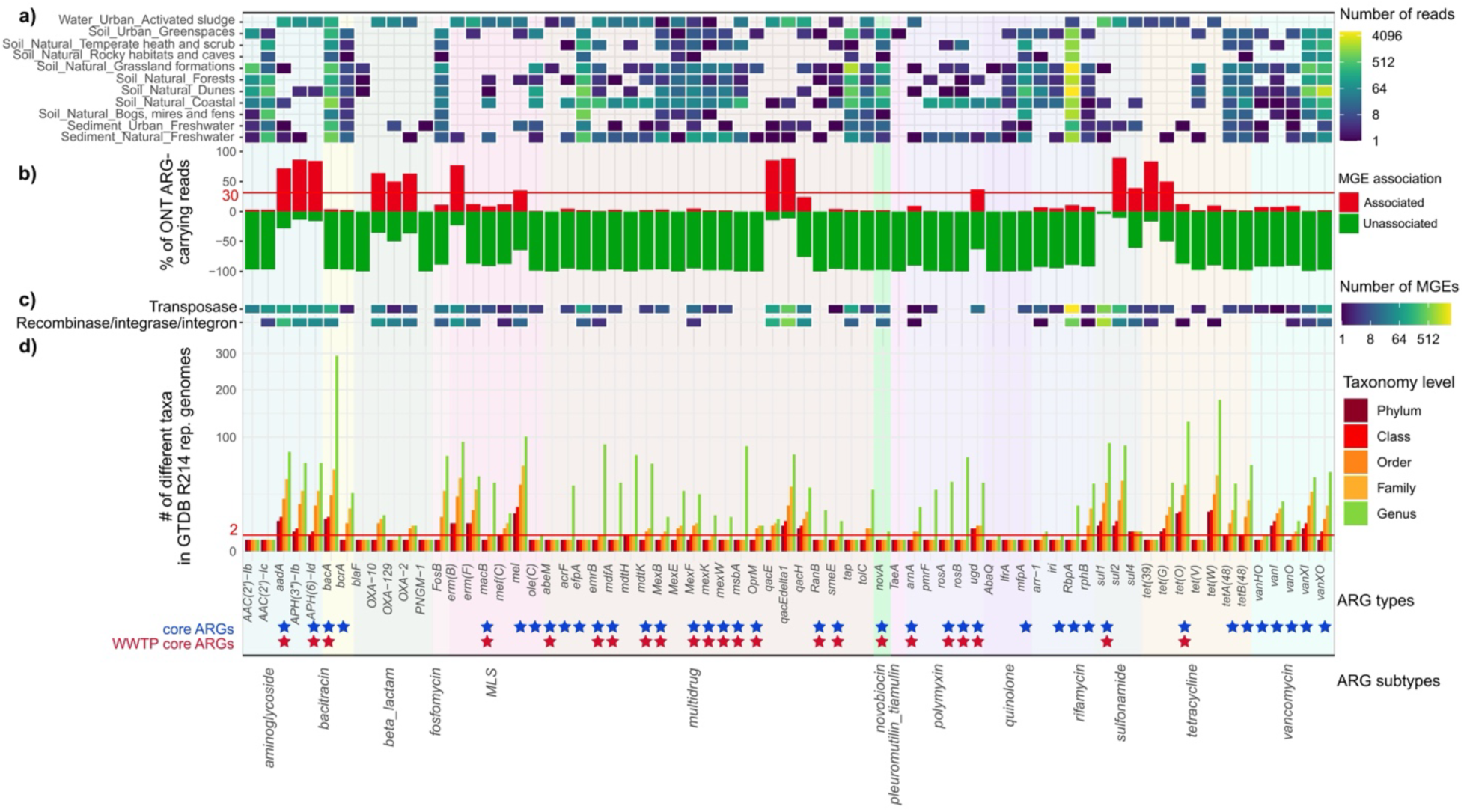
Mobility, host range, and hosts of the commonly (cross-habitat) abundant ARGs. **a)** Heatmap illustrating the specific habitats each ARGs were detected in, colored by the number of long reads carrying the corresponding ARGs. **b)** Diverging barchart showing the proportions of ARG-carrying ONT quality-trimmed reads with (red) and without (green) MGEs identified within 5kb distance to the ARG for those 69/83 MGE-associated commonly abundant ARGs. **c)** Heatmap displaying the close association (within 5kb radius of the ARGs) between the commonly abundant ARGs and MGEs. **d)** Barplot demonstrating the phylogenetic barrier of the 69/83 MGE-associated commonly abundant ARGs in GTDB (R214) representative genomes at different taxonomic levels. MLS: Macrolides-lincosamides-streptogramines.

Generally, we found that stronger associations between ARGs and MGEs indicated broader host ranges (Figure 4d). One set of these commonly abundant ARGs had limited associations to MGEs, but exhibited broad host ranges extending beyond the genus level. This likely stemmed from the long-term microbial evolution of specific ecological microbiota, such as the many vancomycin-ARGs in soil-dwelling vancomycin-producing microbes (Actinomycetes). The presence of these ARGs in soils likely represent the natural baseline resistome. This was confirmed in our Danish MAG data, as many hosts of vancomycin-ARGs were Actinomycetes (SI1 Table S11). Another set of commonly abundant ARGs with limited MGE associations had comparatively narrower host ranges, such as multidrug-ARGs, which appeared to be limited to inter-genera (of the same family)/inter-species (of the same genus) sharing within *Enterobacteriaceae* and *Pseudomonoadaceae* (SI1 Table S11) facilitated by limited MGEs. This is probably linked to the intrinsic (e.g. chromosomally located and vertically inherited) and phylogenetically conserved nature of these ARGs ^67,84^. Although some of them can be plasmid-borne as suggested in CARD (SI1 Table S11), their dissemination may remain constrained by plasmid host range and taxonomic barriers, limiting broad cross-lineage transfer. The set of commonly abundant ARGs with strong MGE association generally had wide host range (above the genus boundary, Figure 4d, SI1 Table S11), such as *aadA*, *erm(B)*, *mel*, *qacEdelta1*, *sul1*, *sul2*, *OXA-2*, *ugd*, *tetO*, and *tetW*. We suggest these ARGs are prioritised for future monitoring investigations, as they are potentially highly mobile across habitats.

### Environmental ARG mobility is shaped by habitats and ARG types, largely non-viral and pathogen-linked

The spread of ARGs within and between One Health sectors (animal, human and environment), is key information needed to tackle current AR challenges. We next investigated HGT potential of the MFD data (assemblies and MAGs from the Nanopore datasets) and compared it to publicly available MAGs derived from human gut, pig gut, marine, and soils, encompassing human, animal, and environmental (global scale) sectors, while MFD data representing the environmental (Danish national-scale) sector, to determine whether ARG dissemination potential varied across habitats and ARG types.

Overall, the environmental sector had lower ARG HGT potential compared to the clinical and animal sectors. As observed in Figure 5a, the incidence of MGEs is always higher in human pathogens than in animal guts and environmental habitats, regardless of the genetic distance from an ARG. This may be partly driven by the representation of human-associated elements in reference databases. Additionally, although soils had the highest proportion of ARG-carrying MAGs (15.2%, 1,923/12,635 MAGs, SI Table S10) and multi-ARG-carrying MAGs (27.6%, 530/1,923), they had the lowest proportion of ARG-carrying MAGs with a nearby MGE (11.6%, compared with 29.0% in AS and 38.3% in sediments, within 5kb radius of ARG). This suggests soils to be reservoirs of ARGs ^85^ where ARGs are largely maintained on the chromosome (Figure 5b) and shaped by long-term microbial evolution, but anthropogenic activity and environmental connectivity select and enrich those that are mobile. Furthermore, WWTPs had greater ARG HGT potential than soils and sediments, suggesting its importance in the stewardship of resistome spread (Figure 5a).

**Figure 5.**
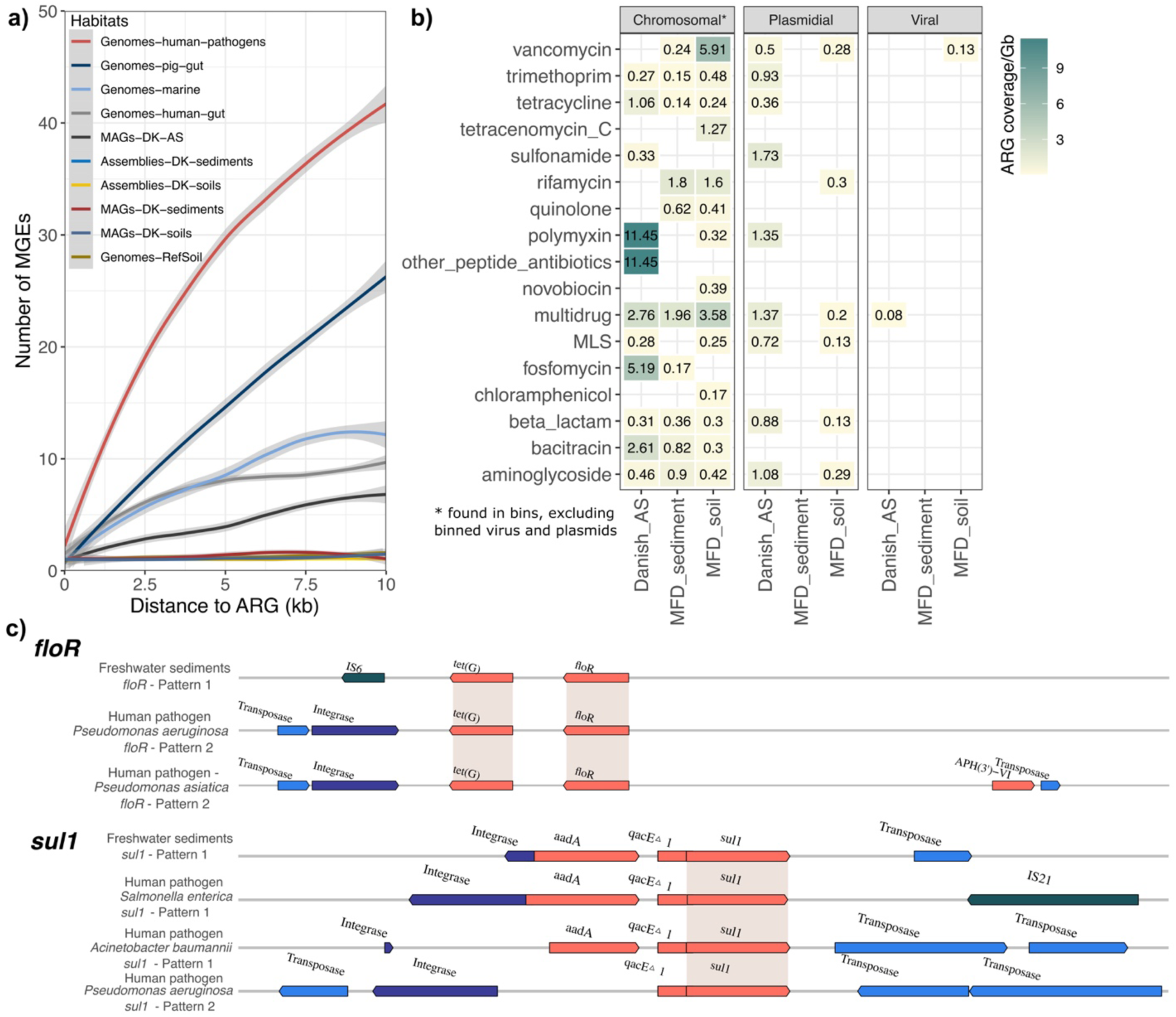
Transfer potential, genetic background, and sharing of ARGs between human pathogens and the Danish environment. **a)** Within-habitat ARG HGT potential measured by the number of MGEs within different distances to ARGs. Counts of transposes, integrases, and recombinases were used as a proxy for the number of MGEs. Human pathogens had significantly higher HGT transfer at all step distances to ARGs (*p* < 0.05, Fisher’s exact test); **b)** Genetic background of ARGs (chromosome, putative plasmids, and putative viruses) shown in heatmap, measured in ARG coverages normalized to per Gb metagenome data for each ARG type based on ARG-carrying contigs, taking contig coverages into consideration; **c)** Two example ARGs (*floR* and *sul1*) that could potentially be shared between environmental ARGs and ARGs carried by human pathogens. Genomic structures of potentially mobile ARG-carrying patterns for highly similar (>98% identity over >98% coverage) ARGs between the environment and human pathogens. Shading denotes the highly-similar ARGs used to align the genomic patterns. DK: Denmark, referring to MFD Nanopore datasets. from MLS: Macrolides-lincosamides-streptogramines.

Interestingly, some ARGs seemed to be differentially mobilizable across ARG types and the environmental habitats in which they reside (Figure 5b). For example, aminoglycoside-, MLS-, and beta-lactam-ARGs appeared to be more plasmid-borne in AS, but more chromosomally located in soils (Figure 5b). Additionally, higher coverages of plasmidial context was observed for sulfonamide-, trimethoprim-, MLS-, beta-lactam-, and aminoglycoside-resisting ARGs in AS than in chromosomal context. This supplements previous observations that these ARGs are more frequently found on plasmids than on chromosomes in AS ^86^. In soils and sediments, ARGs are more typically found on chromosomes than on plasmids.

Although viruses are proposed as a key reservoir for ARG transfer in different studies ^87^, there is no consensus on the contribution of viruses to ARG spread. In this study, viruses were rarely found to encode ARGs, suggesting that they are unlikely to substantially contribute to the spread of ARGs in Danish soils, sediments, and AS. Out of all the 7,042 complete or HQ viral contigs, no viruses contained ARGs (SI2 Table S11a). Expanding the investigation to all 92,036 MQ and low-quality (LQ) viral contigs revealed only 4 potential ARG-carrying LQ viral contigs in Danish AS and soil (Figure 5b). This suggests that viral carriage of ARGs may not be widespread (or may be unstable) in the environment ^88,89^, but may be of selective advantage under human impact ^90^. Our study used bulk metagenomes, which likely limits viral genome recovery compared with virus-enriched viromes ^91^. Given the limited detection of virus-borne ARGs, we note that it is not possible to draw strong conclusions about viral participation in ARG spread across Danish environmental habitats. Future efforts focused on viruses in ARG ecology could use deeper long-read sequencing or virus-enrichment approaches to recover more complete viral communities.

Having identified habitat- and ARG-type-associated linkages to resistome mobility, we next asked whether some environmental mobile ARGs showed links to human pathogens. Despite our observation that the environmental resistome had limited transferability, especially in soils and sediments, we identified 17 mobile environmental ARGs that were highly similar (98% nucleotide identity) to mobile ARGs carried by pathogens (Figure 5c) (SI1 Figure S12). Seven of these ARGs were frequently found next to MGEs in our long read data, i.e., *aadA*, *APH(3’)-Ib*, *erm(B)*, *qacEdelta1*, *sul1*, *sul2*, and *tet(G)*. Additionally, the broad taxonomic host ranges of these ARGs, based on our MAG data (Figure 4d), highlights their potential for transfer between the environment and human pathogens. This potential is further supported in three of the seven ARGs (*aadA*, *qacEdelta1*, and *sul1*), which were frequently associated with class 1 integrons that are strongly implicated in the HGT of ARGs ^92^, especially in pathogens ^93^.

Our results are additionally limited by the use of MAGs to link ARGs to their hosts, which is low-throughout. Although genome-based examination provides confidence in ARG host prediction, host information for ARGs on unbinned contigs are missing, which accounts for 63.8% of ARG-carrying contigs in our data (SI2 Table S11b). Nevertheless, genome-based host tracking is a relatively accurate and cost-effective option that has been shown to agree with HiC-based host tracking ^94^. Additionally, plasmids, an important vehicle of ARG spread, have a huge impact on ARG dissemination ^95,96^, but are largely overlooked in genome-based ARG host-tracking. In our data, for example, 92.6% ARG-carrying plasmidal contigs were unbinned (SI2 Table S11b). Future efforts on HiC ^95^, epicPCR ^97^, and methylation patterns ^98^ will likely expand the knowledge of the microbial hosts and genetic location of ARGs in different habitats.

## Conclusion

Our study provides the first national-scale cross-habitat resistome characterization by surveying more than 7,000 environmental samples, and using both the short-read and long-read sequencing data. Investigating the environmental resistome is an important part of the One Health approach to combating AR. For efficient and cost-effective environmental resistome surveillance, establishing baseline and target ARGs for actionable measures is essential. Here, we assess the distribution and spread of ARGs across Danish habitats, to record the ARGs that represent a consistent, naturally occurring “baseline”. Additionally, the breadth of habitats covered enables investigation of candidate indicator ARGs, permitting ecological source-tracking and ARG “leakage” assessment between urban and natural habitats. By distinguishing baseline and indicator ARGs, our study provides a framework for prioritizing ARG signals relevant to monitoring and stewardship. Additionally, we show that despite the nationally-restricted and reduced antibiotic use in Danish livestock ^99^, Denmark is not free from potential transmission between environmental and pathogen ARGs. This potential transmission is shown by the co-occurrence of MGE-associated highly-similar ARGs between human pathogens and the environment, implying that environmental ARG monitoring is highly relevant.

While our findings reveal habitat-specific patterns in genes associated with AR, we must emphasise that large-scale resistome studies are unable to characterise ARG functions based on homology alone. For example, beta-lactam ARG identification is especially technically limited due to the sharing of functional domains and motifs between ARG and non-ARG sequences ^100^. Future efforts to develop ARG-specific curation criteria for sequence identity and functional confidence, rather than relying solely on homology-based detection in metagenomes, could substantially improve the certainty of ARG identification. In addition, linking ARGs to microbial hosts, MGEs, and transfer potential will further help resolve which environmental resistomes are most likely to contribute to future clinical resistance. Here, we aimed to develop a country-wide profile of all genes that directly and indirectly contribute to AR. This broad foundation is beneficial when planning for future scenarios under the selection pressures of different antibiotics. All in all, our metagenome-based profiling highlights the role of environmental habitats in shaping resistome diversity and dissemination under the One Health framework. Our findings support the idea that diverse microbial habitats can act as ecological barriers to ARG accumulation. Maintaining habitat heterogeneity may therefore help limit resistome homogenisation in human-impacted environments and reduce convergence towards more mobile and clinically relevant ARG pools. We hope this information will spur efforts to support biodiversity in our urban and agricultural landscapes.

## Data availability

The 7,156 MFD short-read Illumina metagenomes and MAGs were collected from the NCBI GenBank under BioProject PRJNA1071982. The 87 Microflora Danica long-read Nanopore metagenomes and recovered MAGs were collected at ENA with BioProject ID: PRJEB58634. The 23 Danish MiDAS Illumina and Oxford Nanopore metagenomes were collected from the NCBI SRA and GenBank databases under the bioproject accession number PRJNA629478. The recovered MQ and HQ MAGs in the Danish MiDAS dataset were collected from Figshare under DOI 10.6084/m9.figshare.c.5277035. All data generated during the study are included as supplementary files.

## Code availability

The methods above indicate the source and versions of the programs and code used for the analyses.

## Acknowledgements

We thank the Microflora Danica Consortium for their invaluable contribution by collecting samples and related metadata across Denmark. Funding was provided by the Poul Due Jensen Foundation, PDJF (grant MicroFlora Danica to M.A. and P.H.N.), Villum Foundation (grant 15510 and 50093 to M.A., grant 13351 to P.H.N., grant VIL60768 to C.M.S), the European Union (ERC grant 101078234 to M.A.). C.M.S. was supported by a Novo Nordisk Foundation Postdoctoral Fellowship grant (NNF20OC0065005).

## Author contributions

Conceptualization: Y.Y., C.M.S., P.H.N., M.A.;

Methodology: Y.Y., L.L., C.L.B., M.S., T.B.N.J.;

Investigation: Y.Y., C.M.S., C.L.B.;

Supervision: C.M.S., M.A., P.H.N.;

Writing of original draft: Y.Y., C.M.S.;

Review and editing: Y.Y., C.M.S., C.L.B., L.L., M.S., T.B.N.J., P.H.N., M.A.

## Supplementary Information

Supplementary Information 1 (SI1)

Supplementary Information 2 (SI2)

Supplementary Information 3 (SI3)

Supplementary Information 4 (SI4)

