## Supplementary material for "Ecological specificity and interconnectivity in Danish environmental resistomes": SI1

### S1. Supplementary methods

In total, over 10,000 samples were collected from various environmental habitats across Denmark through collaborative sampling efforts in the Microflora Danica project (MFD) [1]. For mapping the national resistome profiles, we removed 1) all agricultural soils (due to limited regulatory permissions); 2) samples with low quality-filtered sequencing depths (i.e., less than 2 million trimmed read pairs); 3) samples from habitats of low sample number ( < 20 samples); and 4) samples from enrichment landfill and biogas reactors, resulting in 7,156 samples, as opposed to the 10,000+ sample in the Microflora Danica project. Analysis of the recovery of ARG abundances under different sequencing depths suggested that 2 Mtrps could sufficiently capture highly representative resistome abundances, but there could be much uncovered diversity of low-abundance ARGs at this sequencing depth (SI1 Figure S2-3). An average of 11.29 million trimmed read pairs (Mtrps, ranged from 2.02 to 133.48 Mtrps) per sample was achieved, with averages in soils (n = 5,298), sediments (n = 1,617), and waters (n = 241) being 11.51 Mtrps, 10.73 Mtrps, and 10.17 Mtrps, respectively (SI2 Table S3).

### S2. Workflow overview.

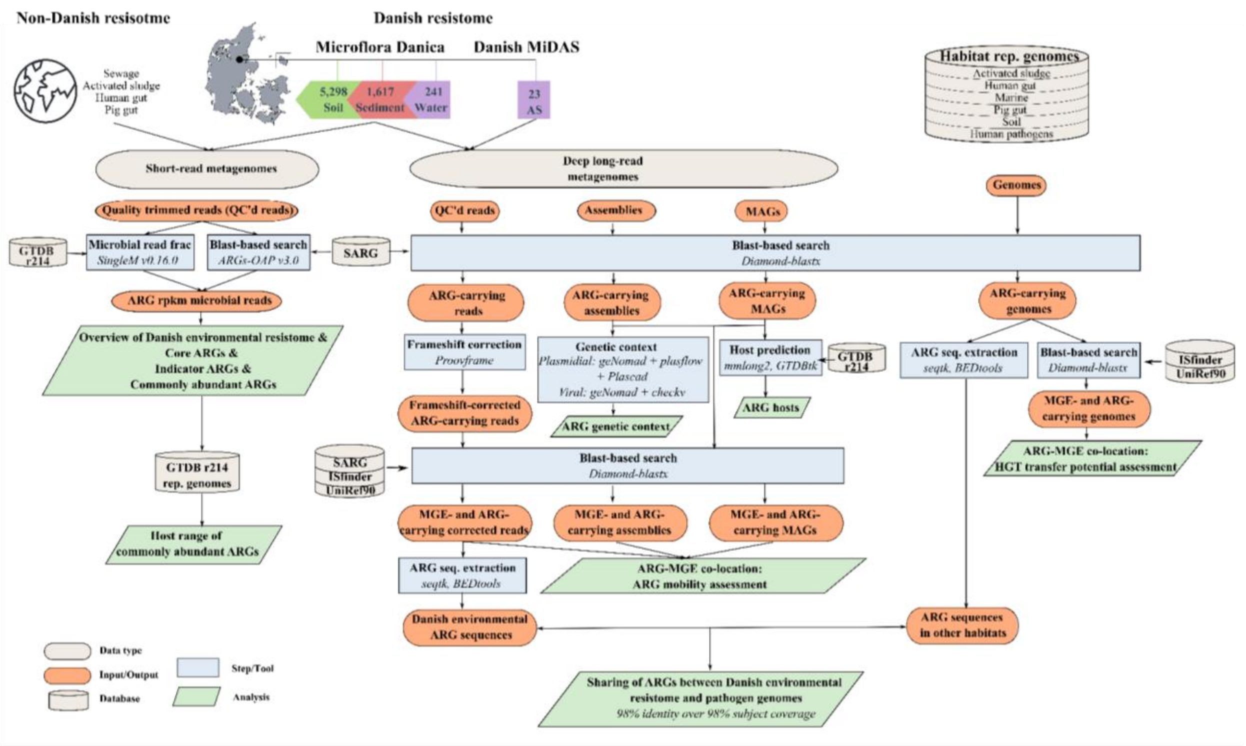

**Figure S1. Overall workflow of this study.**

### S3. Effect of metagenome sequencing depth on the recovery of ARG diversity and abundances.

Four deeply Illumina-sequenced soil samples were used for testing the effect of sequencing depths on the recovery of ARG diversity and abundances. The trimmed sequencing data were first subsampled using seqkit (v2.5.1, https://github.com/shenwei356/seqkit/releases) to 0.1 million read pairs (M_PE), 0.25M_PE, 0.5M_PE, 1M_PE, 2M_PE, 4M_PE, 8M_PE, 16M_PE, 32M_PE. The detection and quantification of ARGs were then performed on these subsampled sequencing datasets and the unsubsampled full datasets (All) using ARGs-OAP (v3.2.4, https://github.com/xinehc/args_oap).

Overall, the total ARG abundances were well-reflected when a minimum of 2 million read pairs (2M_PE) was satisfied when compared to the total ARG abundances estimated from the unsubsampled full datasets (Figure S2a, Figure S3a). Although low-abundance ARG types (Figure S2a) and ARG subtypes (Figure S3a) were to be missed at this sequencing depth, the detected high-abundance ARGs were enough for the recovery of the total ARG abundances. In terms of the estimation of the abundances of each detected ARG types (Figure S2b) and ARG subtypes (Figure S3b), strong spearman correlations ($p < 0.01$) were achieved between the 2 million read pairs (2M_PE) subsampled datasets and the unsubsampled full (All) datasets with spearman $\rho$ 0.87 - 0.94 and $\rho$ 0.57 - 0.66 for ARG types and ARG subtypes, respectively.

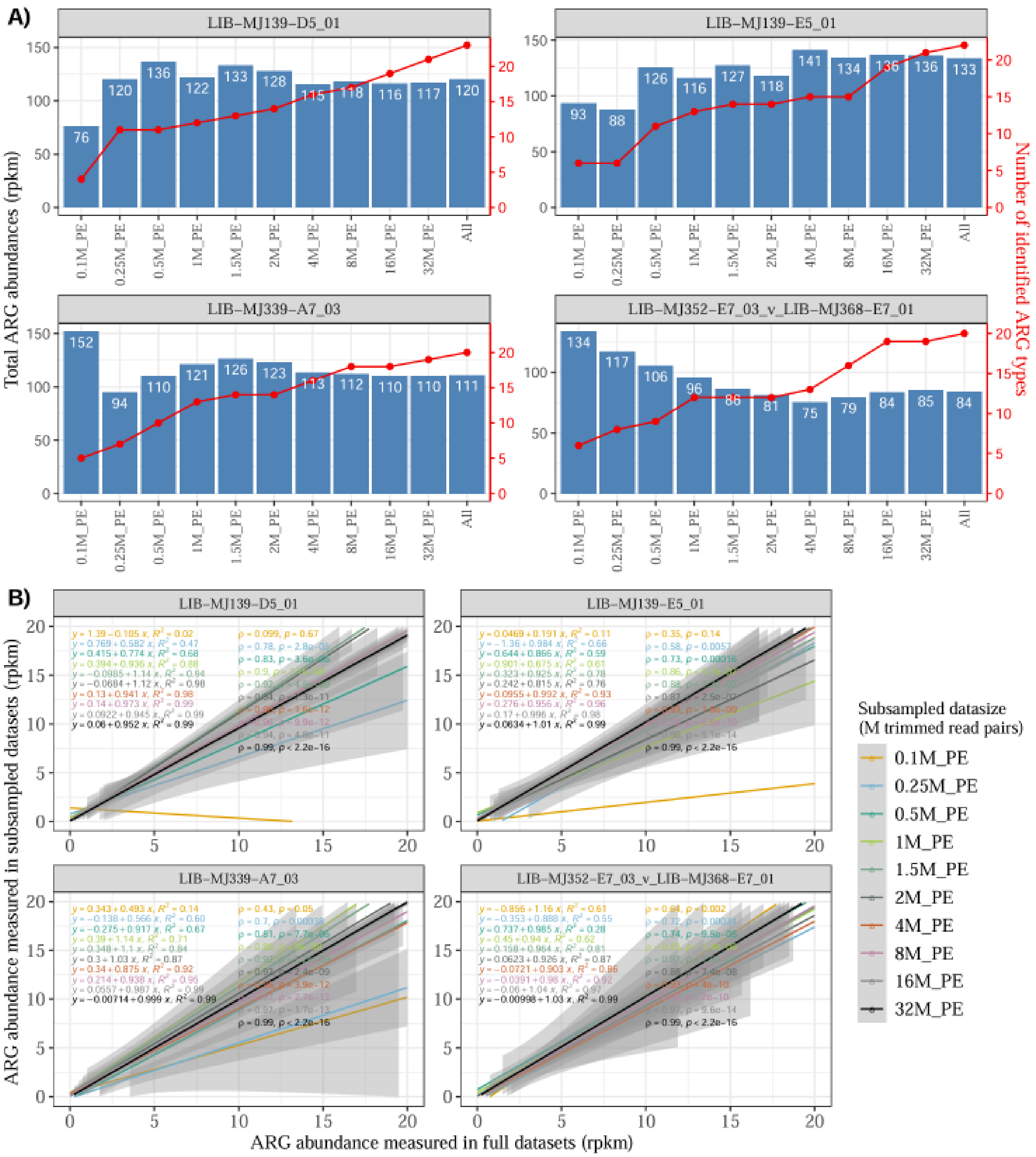

**Figure S2. Recovery of ARG profile at the ARG type level** under different sequencing depths using 4 most-deeply shallow metagenomes. A) A combined barplot and line plot showing the recovery of total ARG abundance and ARG type richness, respectively, under different subsamples. B) Spearman correlation analysis (ρ) and linear regression analysis ($R^2$) showing the agreement on the abundances of each ARG type in rpkm between each subsampled datasets and the unsubsampled full datasets. Line colors represent different sequencing depths of the subsampled datasets: 0.1 million read pairs (M_PE), 0.25M_PE, 0.5M_PE, 1M_PE, 2M_PE, 4M_PE, 8M_PE, 16M_PE, 32M_PE.

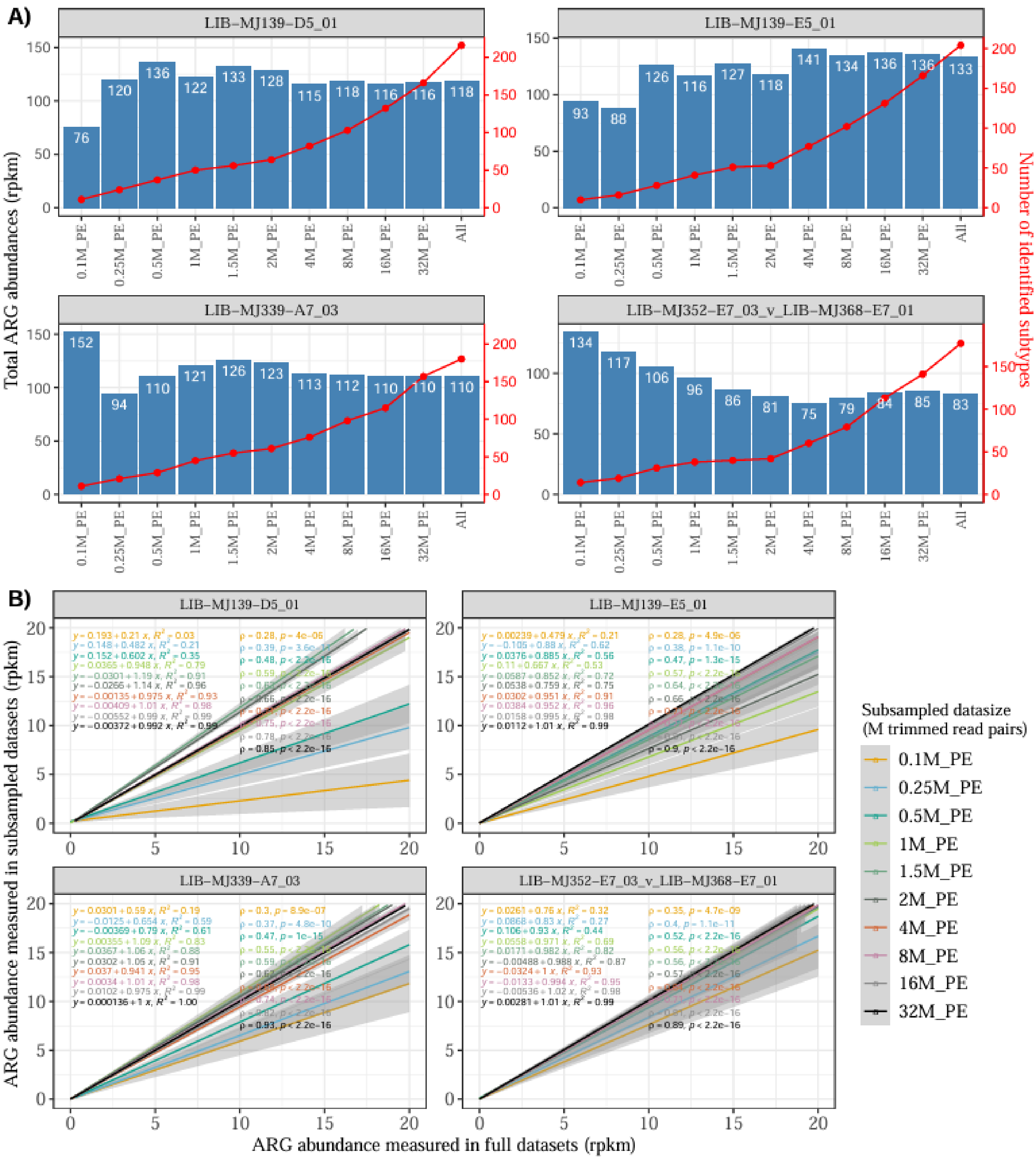

**Figure S3. Recovery of ARG profile at the ARG subtype level** under different sequencing depths using 4 most-deeply shallow metagenomes. A) A combined barplot and line plot showing the recovery of total ARG abundance and ARG subtype richness, respectively, under different subsamples. B) Spearman correlation analysis (ρ) and linear regression analysis ($R^2$) showing the agreement on the abundances of each ARG subtype between each subsampled datasets and the unsubsampled full datasets. Line colors represent different sequencing depths of the subsampled datasets: 0.1 million read pairs (M_PE), 0.25M_PE, 0.5M_PE, 1M_PE, 2M_PE, 4M_PE, 8M_PE, 16M_PE, 32M_PE.

### S4. Total ARG abundances per sample across different habitat types.

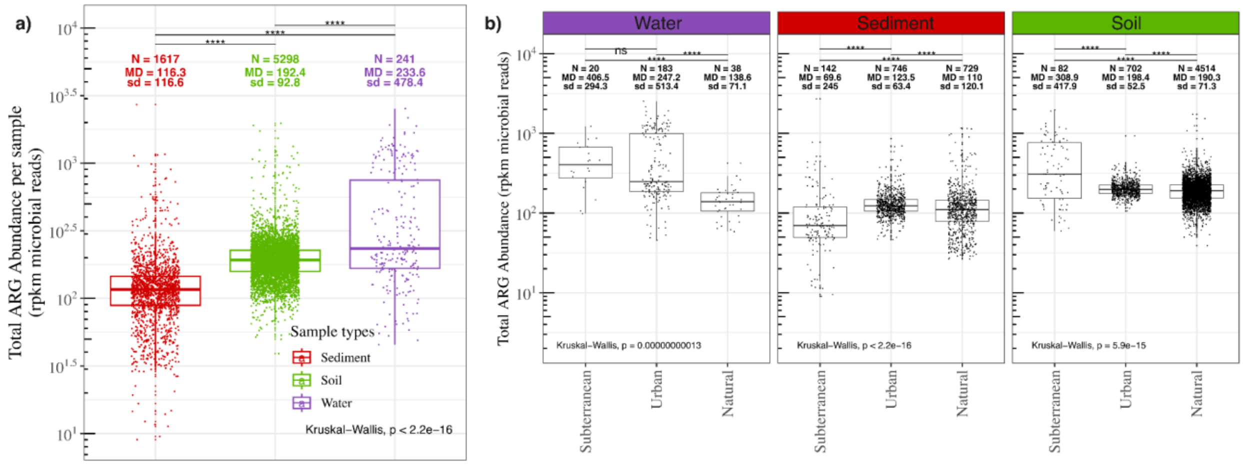

**Figure S4. Total ARG abundances (rpkm microbial reads) per sample** stratified to different a) sample types; b) area types within each sample type. Differences between multiple area type groups within each sample type were tested by Kruskal-Wallis test. Differences between each of the 2 area types were tested by Mann-Whitney test. Asterisks and “ns” indicate statistical significance of differences between area types: ns: $p > 0.05$, *: $p \leq 0.05$, **: $p \leq 0.01$, ***: $p \leq 0.001$, and ****: $p \leq 0.0001$. MD: median, N: sample number, sd: standard deviation.

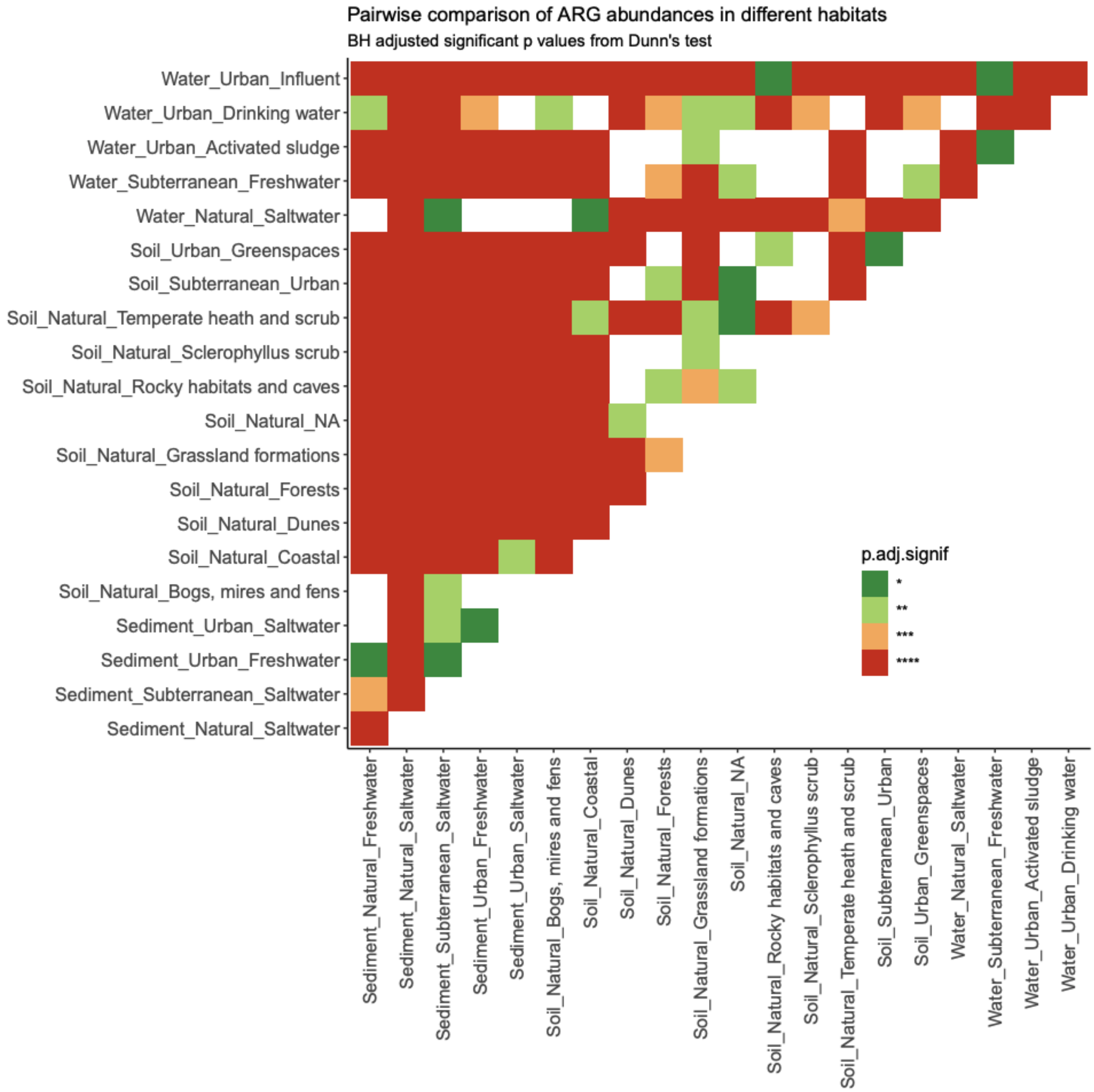

**Figure S5. Pairwise comparison of ARG abundances** by the *post-hoc* Dunn's test based on the resistome from different habitats. Color gradation shows the BH-adjusted *p*-values (high: red, green: low). Asterisks and blank indicate statistical significance of differences between habitats: blank cell: $p > 0.05$, *: $p \leq 0.05$, **: $p \leq 0.01$, ***: $p \leq 0.001$, and ****: $p \leq 0.0001$.

### S5. Danish ARG abundances in WWTPs compared to the non-Danish WWTPs.

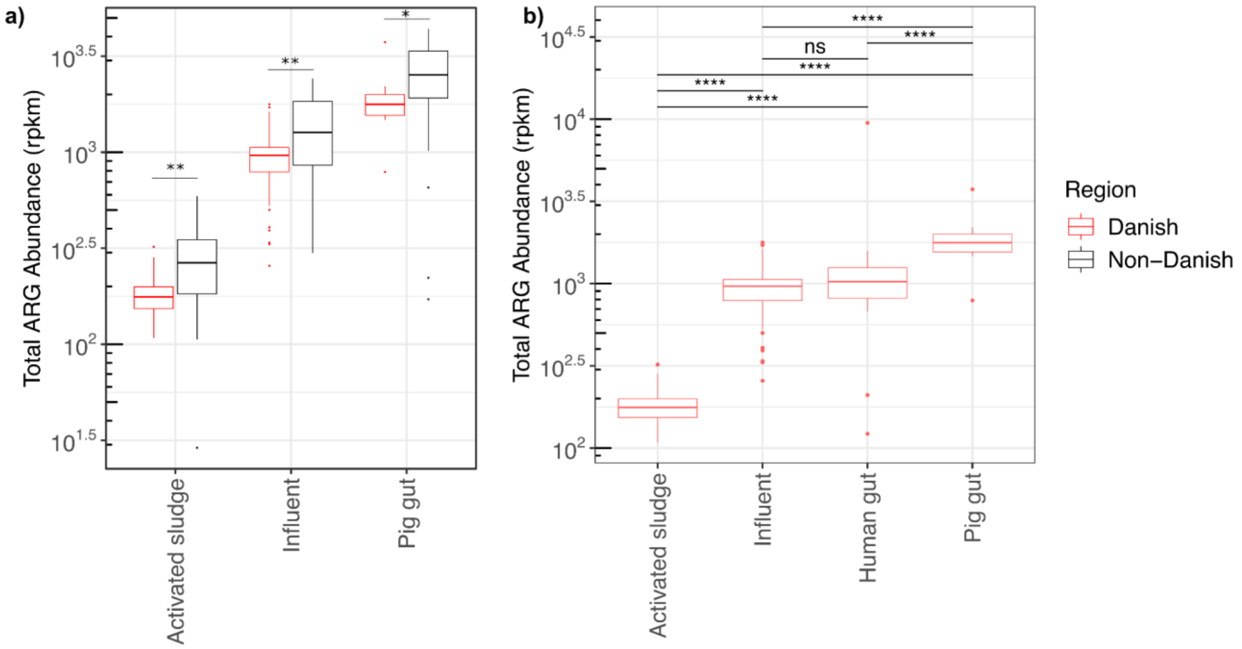

**Figure S6**. Boxplots comparing the total ARG abundances in **a)** Danish and non-Danish WWTP and pig gut samples. ANOSIM based on ARG abundance profiles showed that between-source dissimilarities were substantially greater than within-source dissimilarities (R = 0.809, P = 0.001); **b)** Danish samples in the One Health framework. ANOSIM based on ARG abundance profiles showed that between-source dissimilarities were substantially greater than within-source dissimilarities (R = 0.913, P = 0.001). Wilcoxon test for comparing differences with BH-adjusted *p*-values.

### S6. Ordination plots for resistome profiles of spatially thinned datasets.

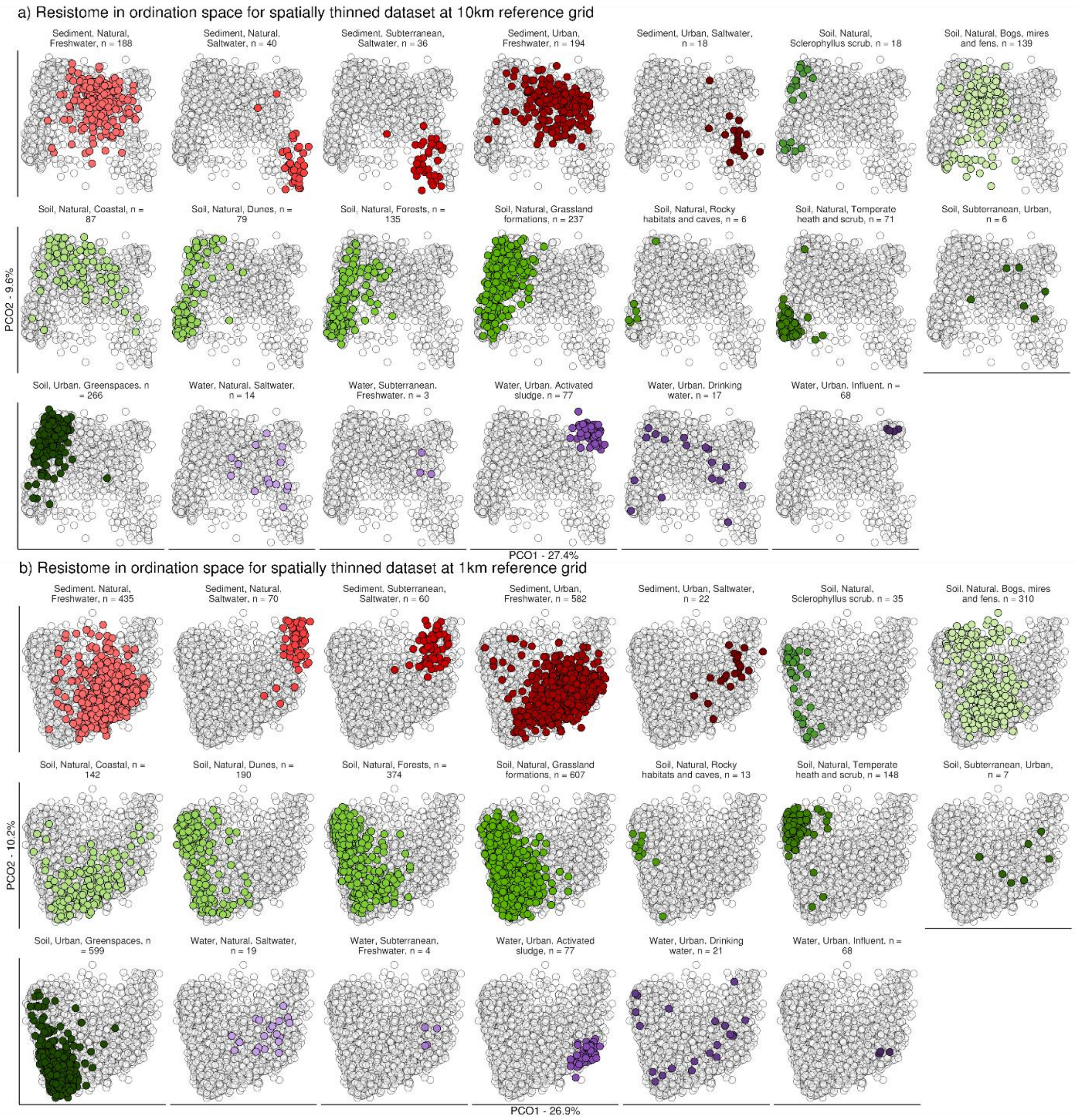

**Figure S7**. PCoA analysis of the resistomes across different environmental habitats based on Bray-Curtis dissimilarity on Hellinger-transformed relative abundances of ARG subtypes for spatially thinned datasets using **a)** the 10-km reference grid of Denmark; and **b)** the 1-km reference grid of Denmark.

### S7. Procrustes analysis to evaluate concordance between microbial composition and resistome composition.

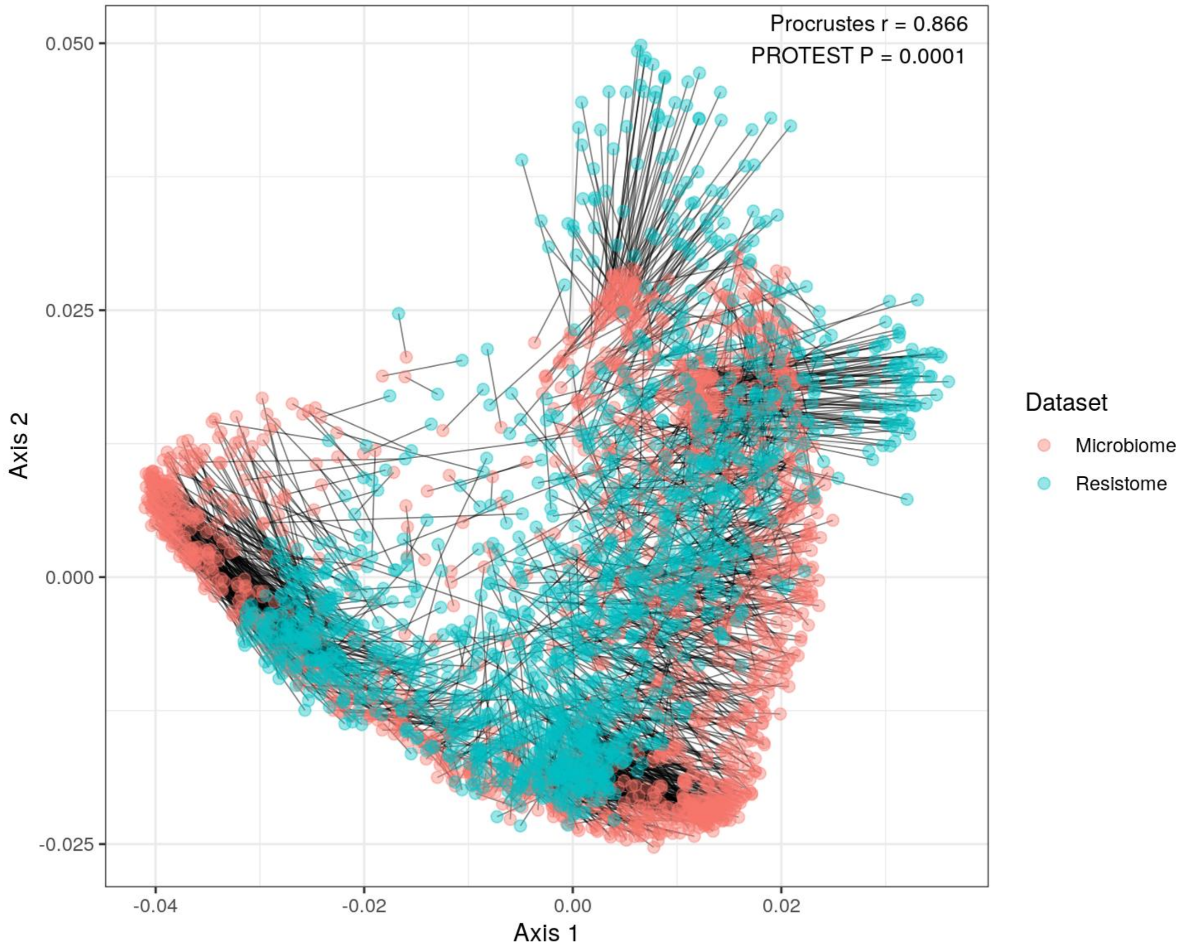

**Figure S8**. Procrustes analysis of the association between resistome composition and microbial community structure for spatially thinned datasets using the 10-km reference grid of Denmark. P-value was generated with a permutational test.

### S8. Detection frequency of identified ARG subtypes across 21 habitats.

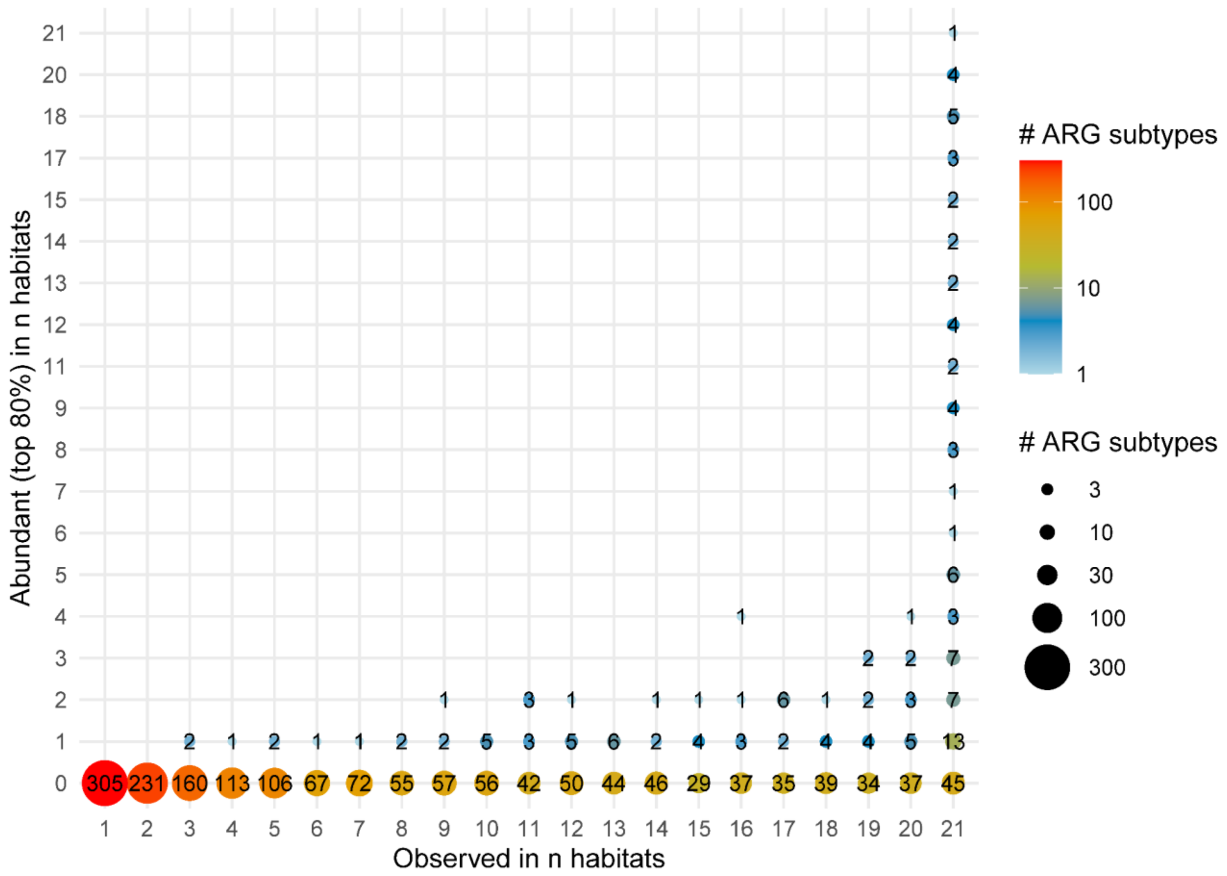

**Figure S9**. Detection frequency of identified 1810 ARG subtypes across 21 different habitats.

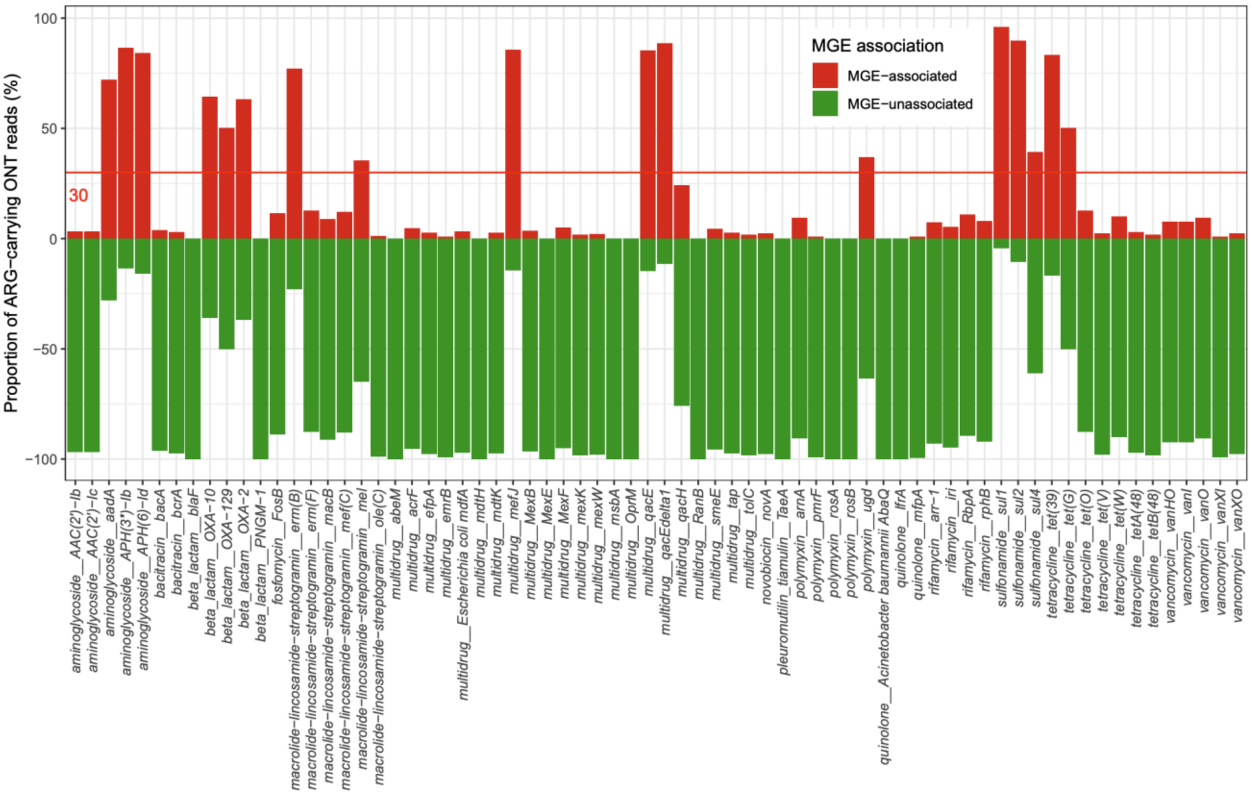

**Figure S10**. Proportion of ARG-carrying quality-trimmed ONT reads having MGEs in the vicinity (with 5 kb radius) of all the detected commonly abundant ARGs in quality-trimmed ONT reads (70/83).

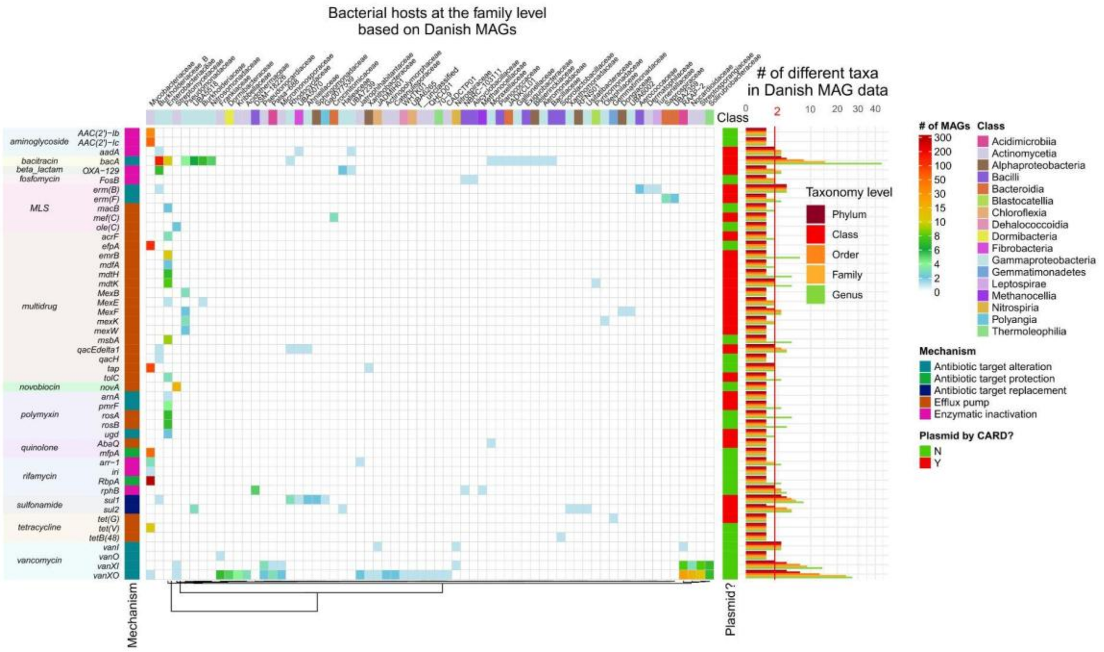

**Figure S11**. The left panel is a heatmap showing the bacterial host ranges at the phylum level for the 48/83 commonly abundant ARGs detected on Danish MAGs. Colors indicate the number MAGs detected with the corresponding ARG-host relationship. Phyla were hierarchically clustered on Euclidean distance using the complete linkage method based on the detection frequency of the ARG-host pairs for different ARGs. The right panel is a barplot demonstrating the phylogenetic barrier of the 48/83 commonly abundant ARGs on Danish MAGs at different taxonomic levels. MLS: Macrolides-lincosamides-streptogramines.

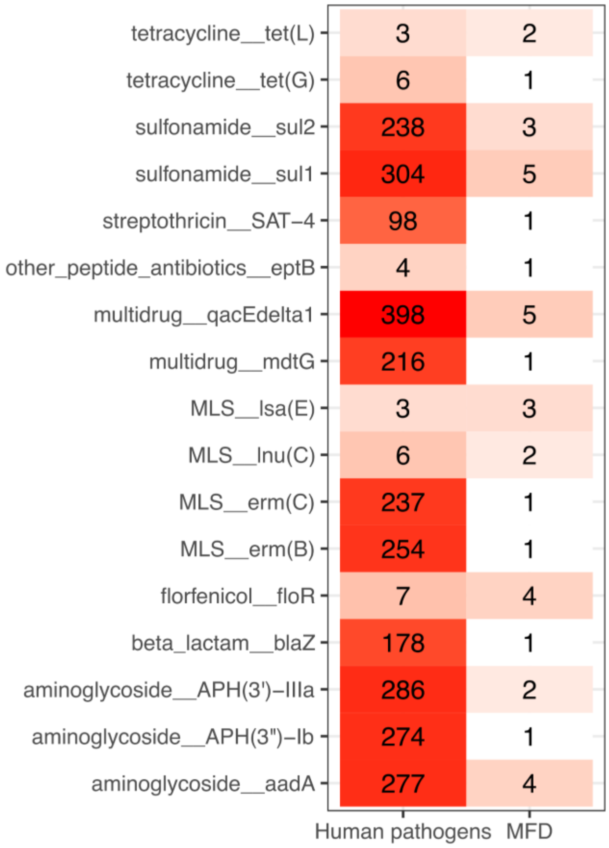

**Figure S12.** Number of hits of highly similar ARGs (98% similarity and 98% coverage over the ARGs on the pathogen genomes) between MFD ONT raw reads and human pathogen genomes.
