## Supplementary material for "Ecological specificity and interconnectivity in Danish environmental resistomes": SI4

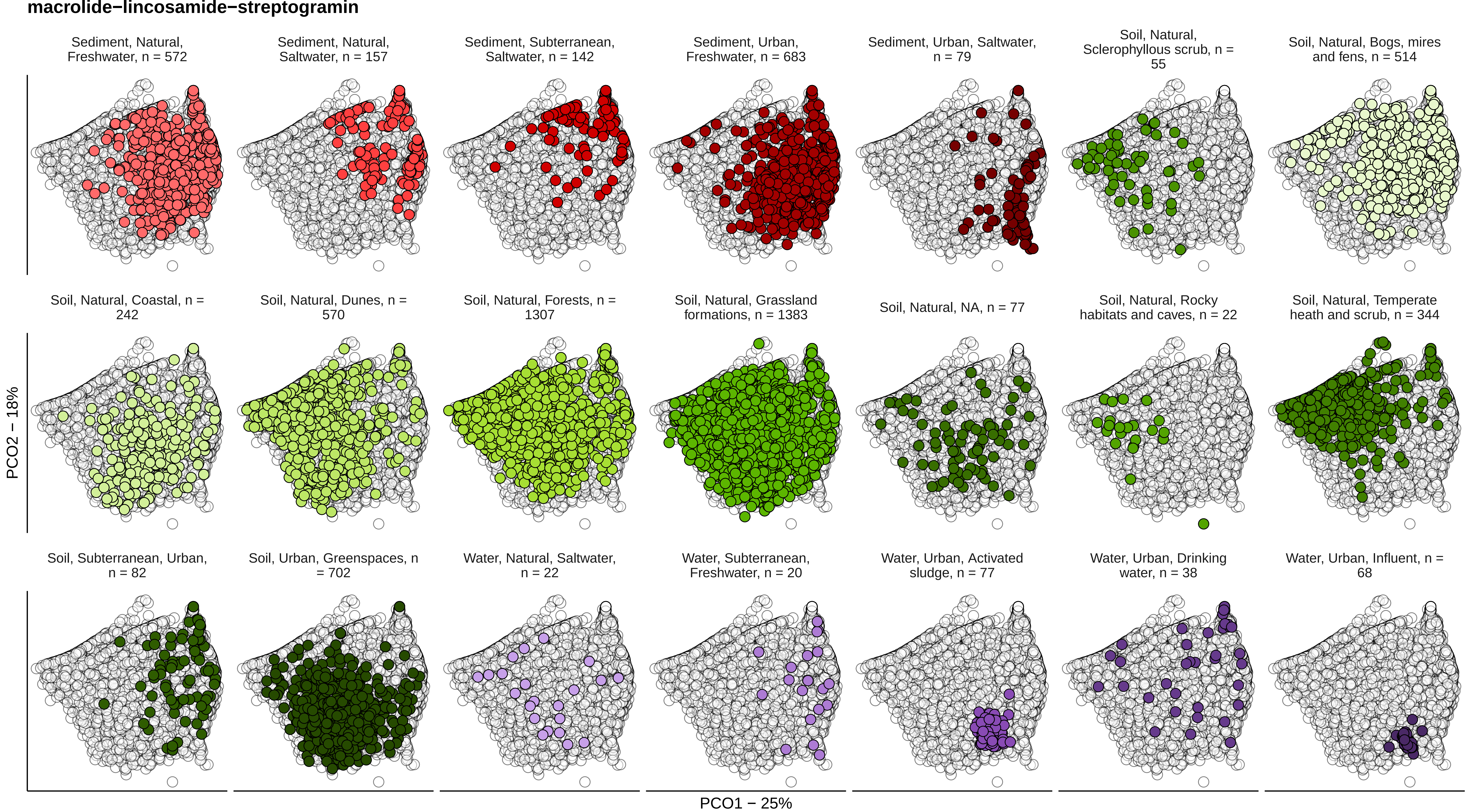

macrolide−lincosamide−streptogramin
Sediment, Natural, Freshwater, n = 572
Sediment, Natural, Saltwater, n = 157
Sediment, Subterranean, Saltwater, n = 142
Sediment, Urban, Freshwater, n = 683
Sediment, Urban, Saltwater, n = 79
Soil, Natural, Sclerophyllous scrub, n = 55
Soil, Natural, Bogs, mires and fens, n = 514
Soil, Natural, Coastal, n = 242
Soil, Natural, Dunes, n = 570
Soil, Natural, Forests, n = 1307
Soil, Natural, Grassland formations, n = 1383
Soil, Natural, NA, n = 77
Soil, Natural, Rocky habitats and caves, n = 22
Soil, Natural, Temperate heath and scrub, n = 344
Soil, Subterranean, Urban, n = 82
Soil, Urban, Greenspaces, n = 702
Water, Natural, Saltwater, n = 22
Water, Subterranean, Freshwater, n = 20
Water, Urban, Activated sludge, n = 77
Water, Urban, Drinking water, n = 38
Water, Urban, Influent, n = 68
PCO2 − 18%
PCO1 − 25%

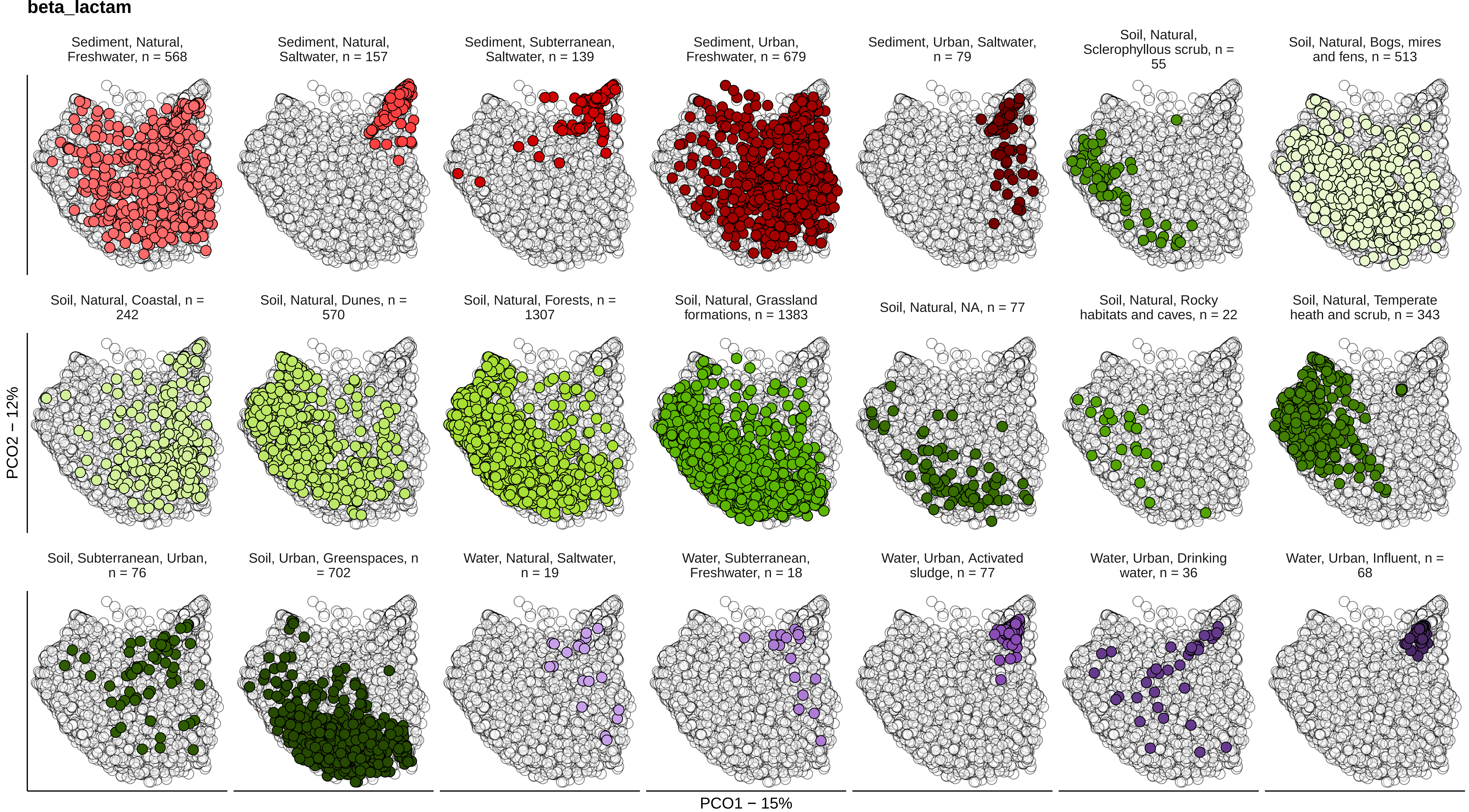
beta_lactam
Sediment, Natural, Freshwater, n = 568
Sediment, Natural, Saltwater, n = 157
Sediment, Subterranean, Saltwater, n = 139
Sediment, Urban, Freshwater, n = 679
Sediment, Urban, Saltwater, n = 79
Soil, Natural, Sclerophyllous scrub, n = 55
Soil, Natural, Bogs, mires and fens, n = 513
Soil, Natural, Coastal, n = 242
Soil, Natural, Dunes, n = 570
Soil, Natural, Forests, n = 1307
Soil, Natural, Grassland formations, n = 1383
Soil, Natural, NA, n = 77
Soil, Natural, Rocky habitats and caves, n = 22
Soil, Natural, Temperate heath and scrub, n = 343
Soil, Subterranean, Urban, n = 76
Soil, Urban, Greenspaces, n = 702
Water, Natural, Saltwater, n = 19
Water, Subterranean, Freshwater, n = 18
Water, Urban, Activated sludge, n = 77
Water, Urban, Drinking water, n = 36
Water, Urban, Influent, n = 68
PCO2 − 12%
PCO1 − 15%

### aminoglycoside

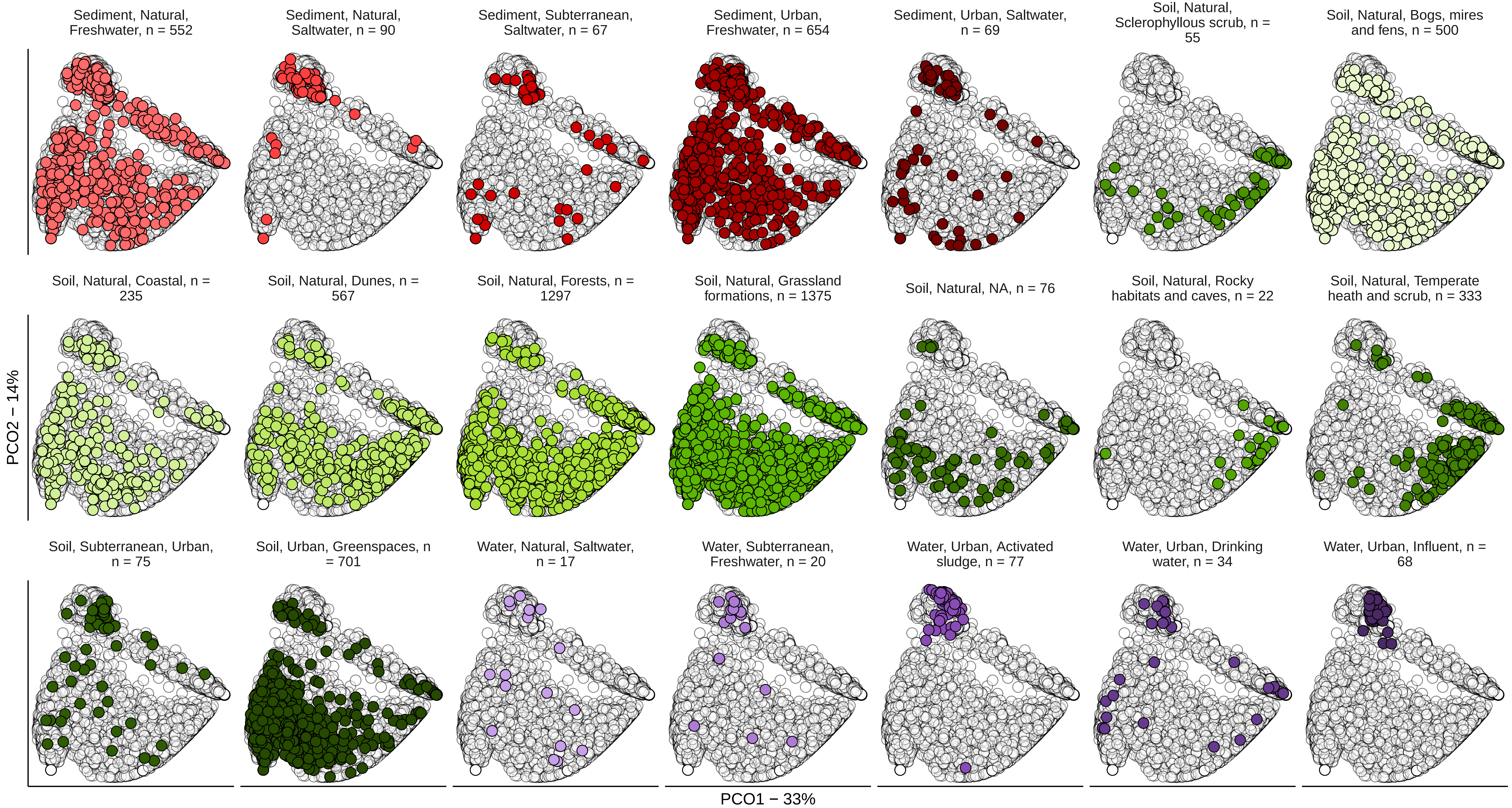

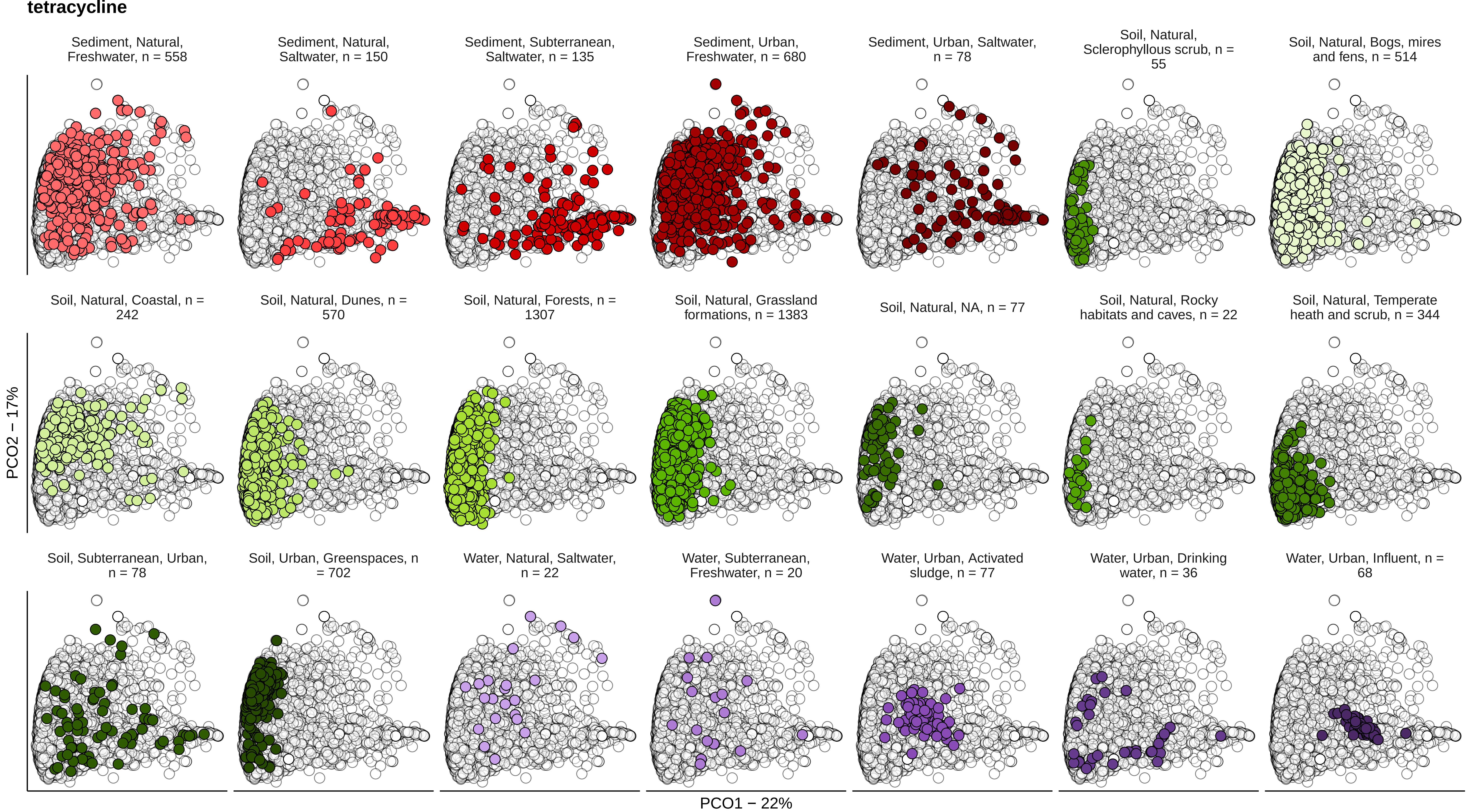
tetracycline
Sediment, Natural, Freshwater, n = 558
Sediment, Natural, Saltwater, n = 150
Sediment, Subterranean, Saltwater, n = 135
Sediment, Urban, Freshwater, n = 680
Sediment, Urban, Saltwater, n = 78
Soil, Natural, Sclerophyllous scrub, n = 55
Soil, Natural, Bogs, mires and fens, n = 514
Soil, Natural, Coastal, n = 242
Soil, Natural, Dunes, n = 570
Soil, Natural, Forests, n = 1307
Soil, Natural, Grassland formations, n = 1383
Soil, Natural, NA, n = 77
Soil, Natural, Rocky habitats and caves, n = 22
Soil, Natural, Temperate heath and scrub, n = 344
Soil, Subterranean, Urban, n = 78
Soil, Urban, Greenspaces, n = 702
Water, Natural, Saltwater, n = 22
Water, Subterranean, Freshwater, n = 20
Water, Urban, Activated sludge, n = 77
Water, Urban, Drinking water, n = 36
Water, Urban, Influent, n = 68
PCO2 − 17%
PCO1 − 22%

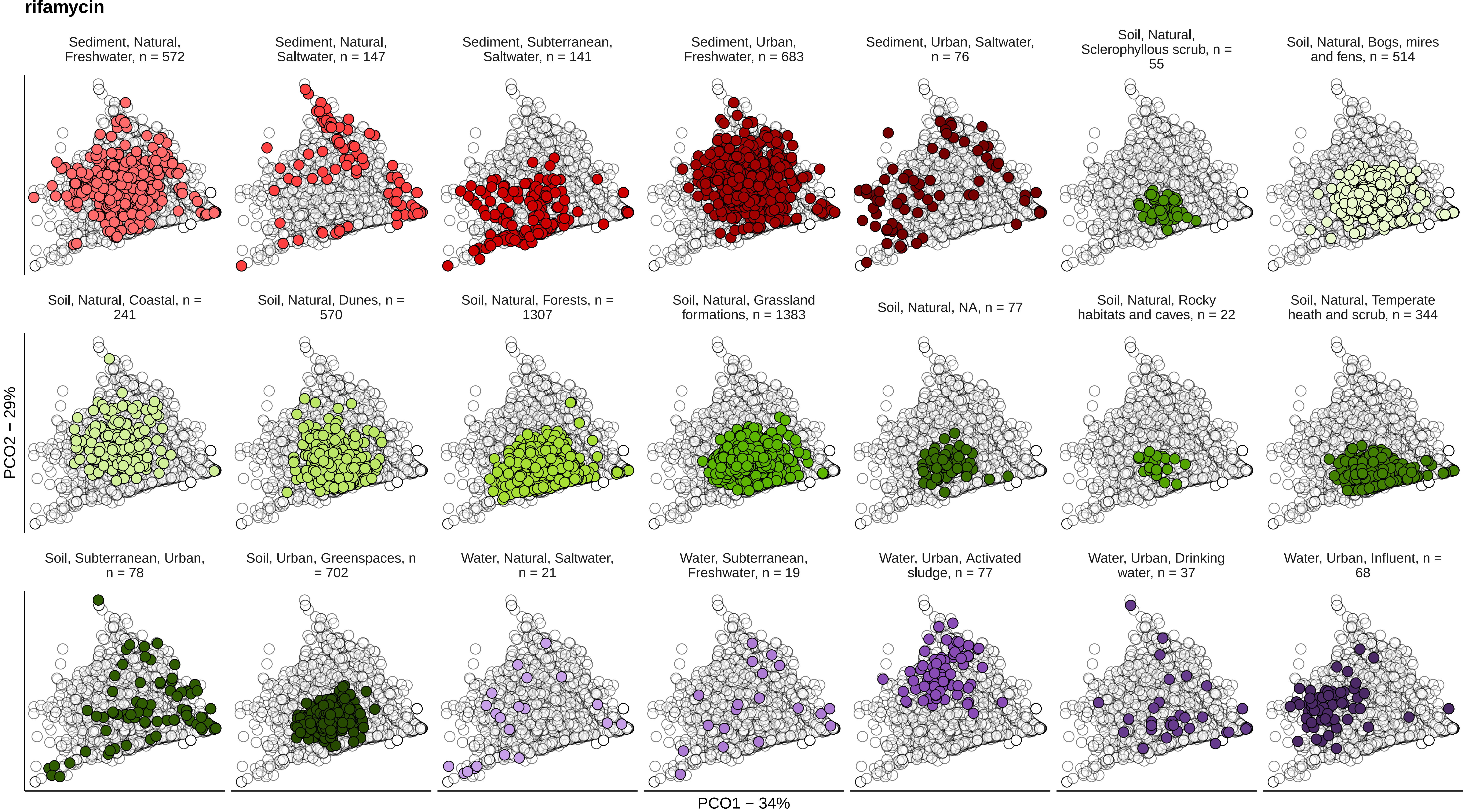

rifamycin
Sediment, Natural, Freshwater, n = 572
Sediment, Natural, Saltwater, n = 147
Sediment, Subterranean, Saltwater, n = 141
Sediment, Urban, Freshwater, n = 683
Sediment, Urban, Saltwater, n = 76
Soil, Natural, Sclerophyllous scrub, n = 55
Soil, Natural, Bogs, mires and fens, n = 514
Soil, Natural, Coastal, n = 241
Soil, Natural, Dunes, n = 570
Soil, Natural, Forests, n = 1307
Soil, Natural, Grassland formations, n = 1383
Soil, Natural, NA, n = 77
Soil, Natural, Rocky habitats and caves, n = 22
Soil, Natural, Temperate heath and scrub, n = 344
Soil, Subterranean, Urban, n = 78
Soil, Urban, Greenspaces, n = 702
Water, Natural, Saltwater, n = 21
Water, Subterranean, Freshwater, n = 19
Water, Urban, Activated sludge, n = 77
Water, Urban, Drinking water, n = 37
Water, Urban, Influent, n = 68
PCO2 − 29%
PCO1 − 34%

**sulfonamide**

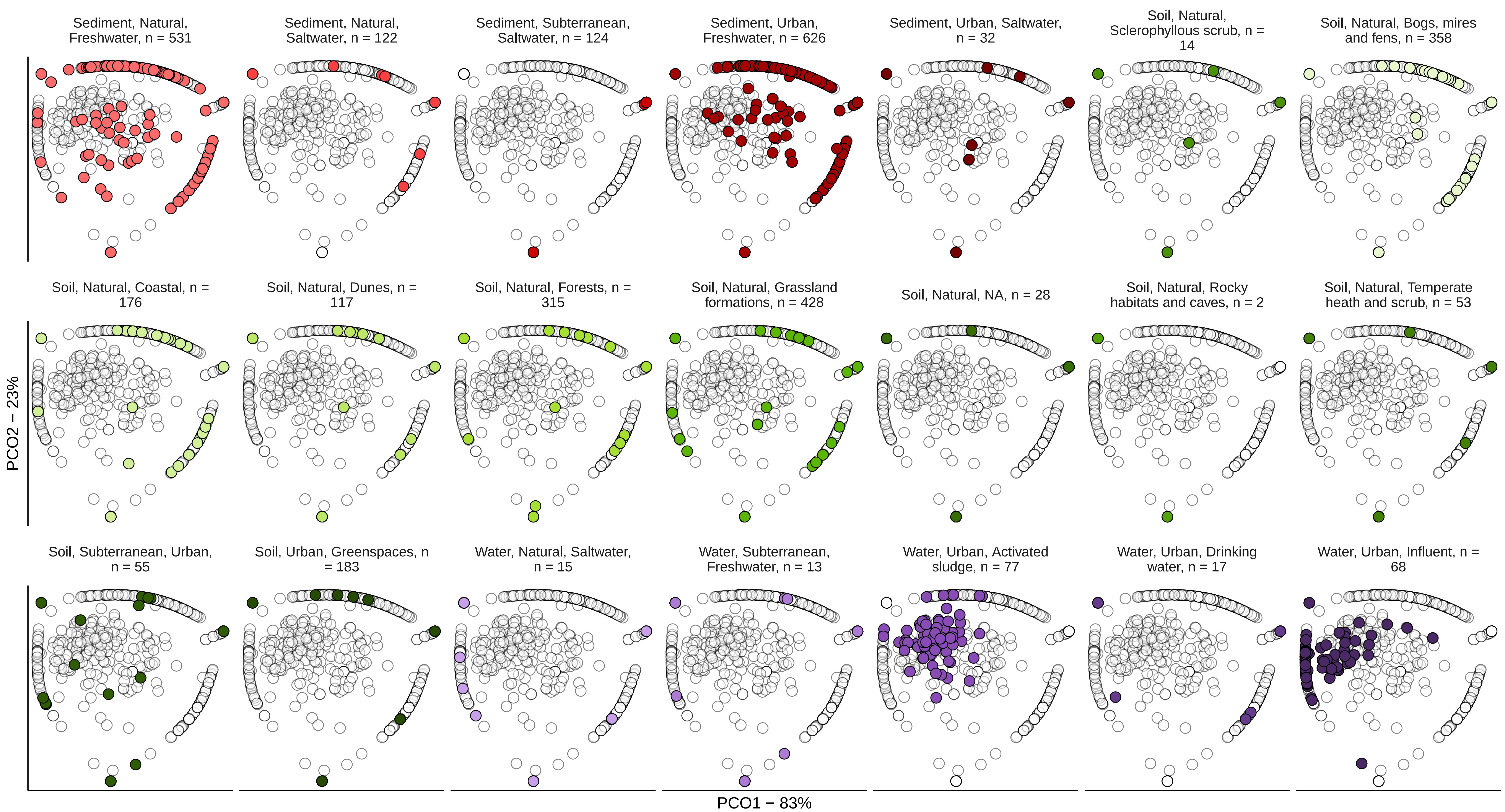

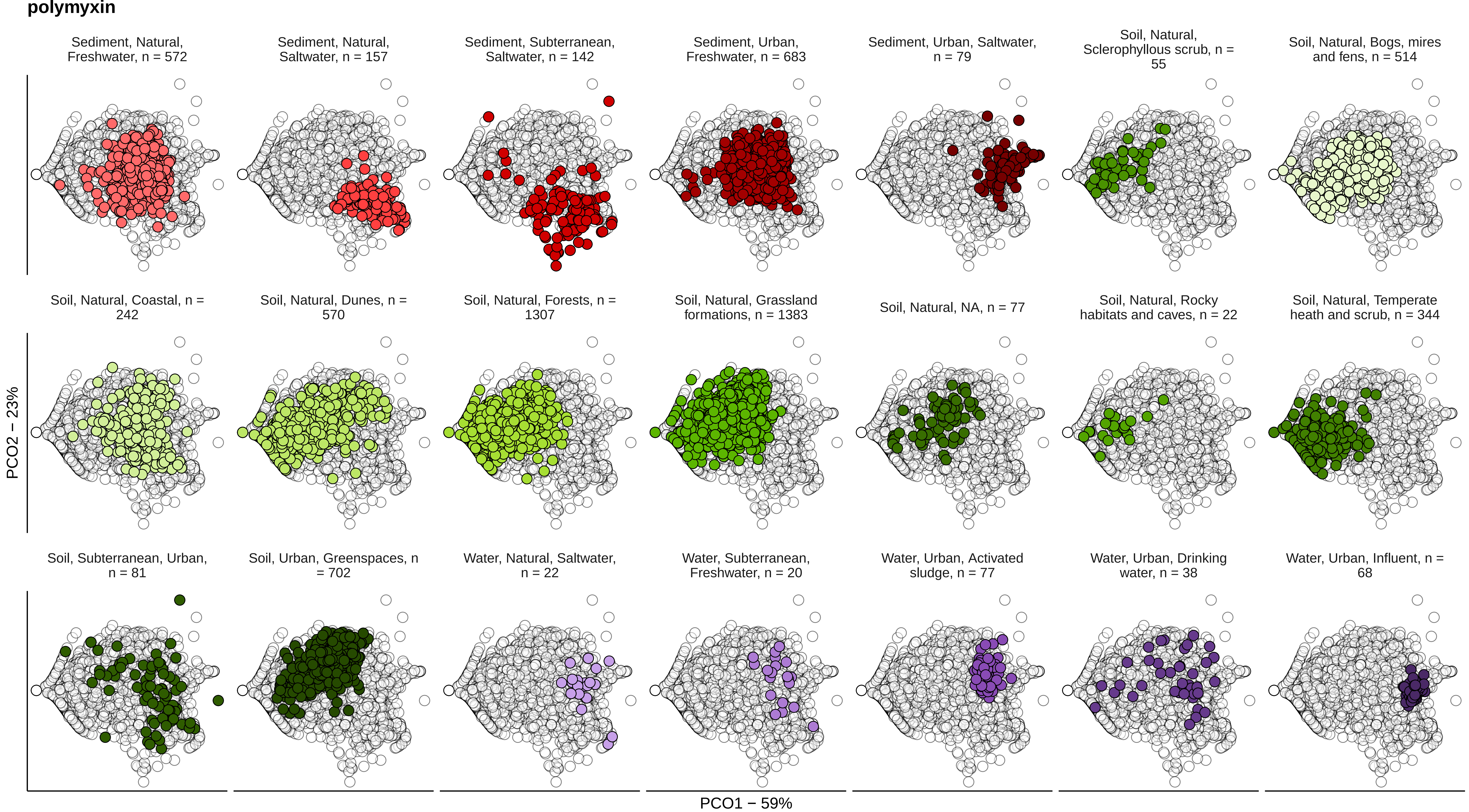
polymyxin
Sediment, Natural, Freshwater, n = 572
Sediment, Natural, Saltwater, n = 157
Sediment, Subterranean, Saltwater, n = 142
Sediment, Urban, Freshwater, n = 683
Sediment, Urban, Saltwater, n = 79
Soil, Natural, Sclerophyllous scrub, n = 55
Soil, Natural, Bogs, mires and fens, n = 514
Soil, Natural, Coastal, n = 242
Soil, Natural, Dunes, n = 570
Soil, Natural, Forests, n = 1307
Soil, Natural, Grassland formations, n = 1383
Soil, Natural, NA, n = 77
Soil, Natural, Rocky habitats and caves, n = 22
Soil, Natural, Temperate heath and scrub, n = 344
Soil, Subterranean, Urban, n = 81
Soil, Urban, Greenspaces, n = 702
Water, Natural, Saltwater, n = 22
Water, Subterranean, Freshwater, n = 20
Water, Urban, Activated sludge, n = 77
Water, Urban, Drinking water, n = 38
Water, Urban, Influent, n = 68
PCO2 − 23%
PCO1 − 59%

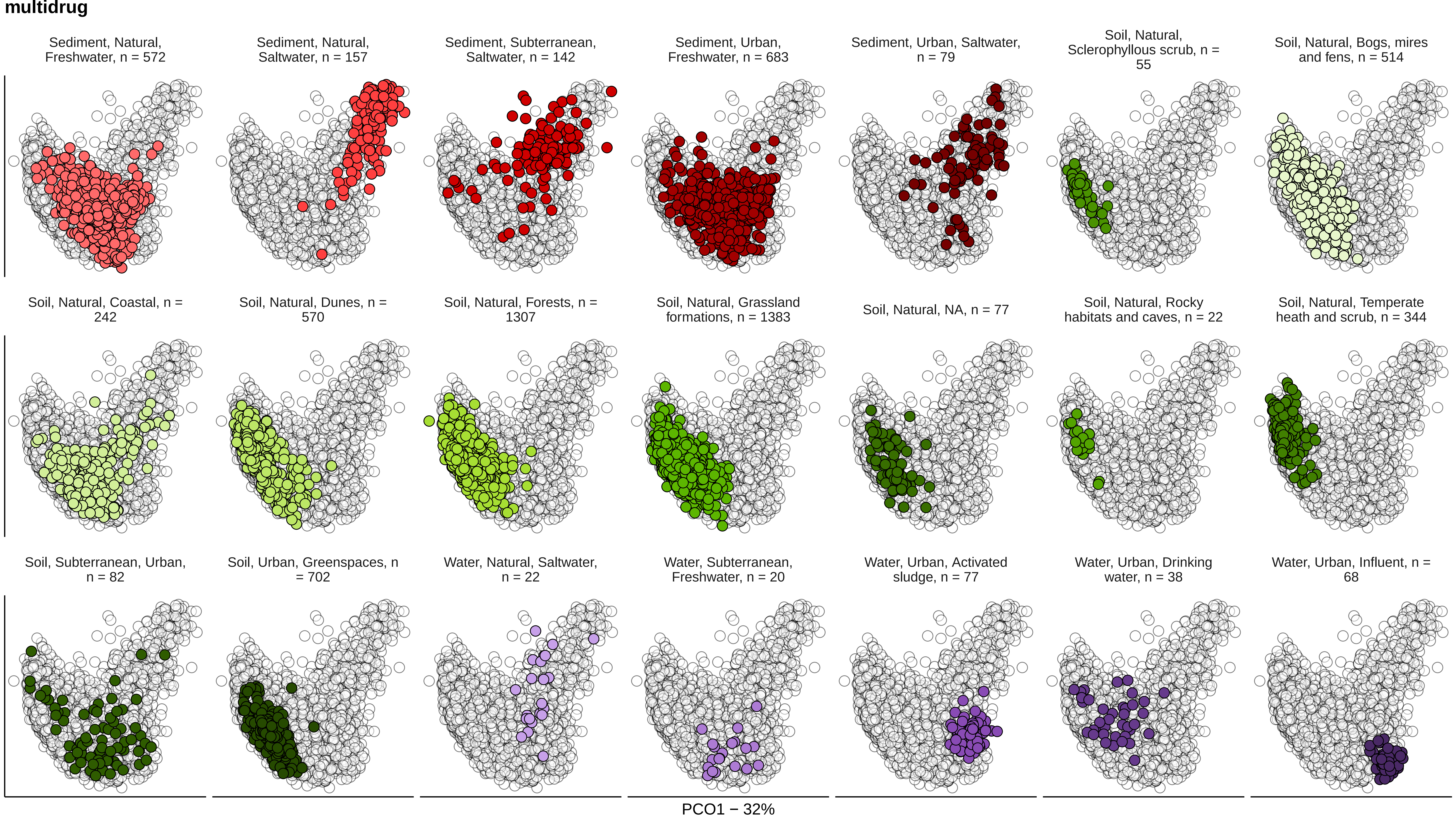

multidrug
Sediment, Natural, Freshwater, n = 572
Sediment, Natural, Saltwater, n = 157
Sediment, Subterranean, Saltwater, n = 142
Sediment, Urban, Freshwater, n = 683
Sediment, Urban, Saltwater, n = 79
Soil, Natural, Sclerophyllous scrub, n = 55
Soil, Natural, Bogs, mires and fens, n = 514
Soil, Natural, Coastal, n = 242
Soil, Natural, Dunes, n = 570
Soil, Natural, Forests, n = 1307
Soil, Natural, Grassland formations, n = 1383
Soil, Natural, NA, n = 77
Soil, Natural, Rocky habitats and caves, n = 22
Soil, Natural, Temperate heath and scrub, n = 344
Soil, Subterranean, Urban, n = 82
Soil, Urban, Greenspaces, n = 702
Water, Natural, Saltwater, n = 22
Water, Subterranean, Freshwater, n = 20
Water, Urban, Activated sludge, n = 77
Water, Urban, Drinking water, n = 38
Water, Urban, Influent, n = 68
PCO2 – 13%
PCO1 – 32%

**vancomycin**

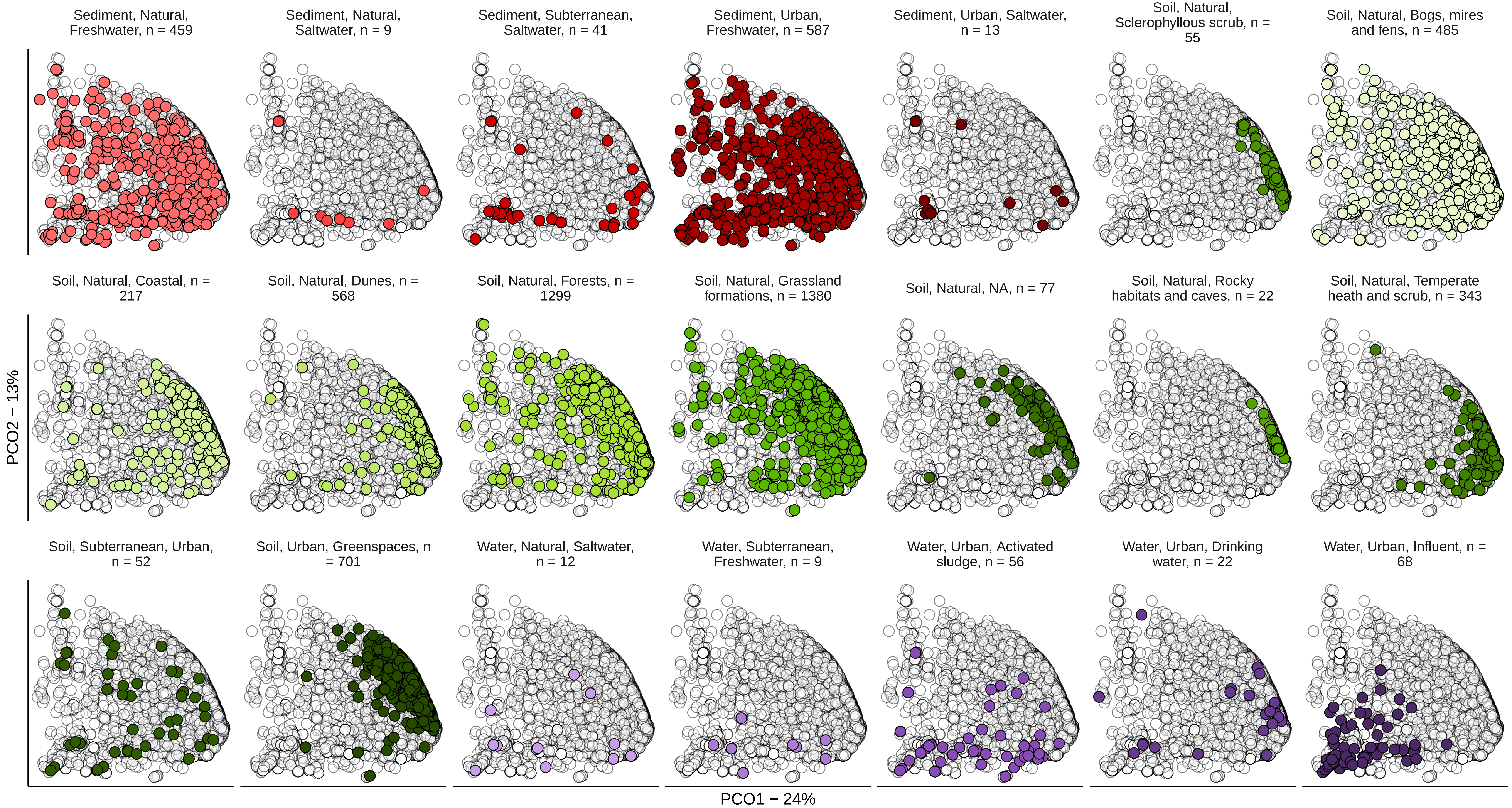
Sediment, Natural, Freshwater, n = 459
Sediment, Natural, Saltwater, n = 9
Sediment, Subterranean, Saltwater, n = 41
Sediment, Urban, Freshwater, n = 587
Sediment, Urban, Saltwater, n = 13
Soil, Natural, Sclerophyllous scrub, n = 55
Soil, Natural, Bogs, mires and fens, n = 485
Soil, Natural, Coastal, n = 217
Soil, Natural, Dunes, n = 568
Soil, Natural, Forests, n = 1299
Soil, Natural, Grassland formations, n = 1380
Soil, Natural, NA, n = 77
Soil, Natural, Rocky habitats and caves, n = 22
Soil, Natural, Temperate heath and scrub, n = 343
Soil, Subterranean, Urban, n = 52
Soil, Urban, Greenspaces, n = 701
Water, Natural, Saltwater, n = 12
Water, Subterranean, Freshwater, n = 9
Water, Urban, Activated sludge, n = 56
Water, Urban, Drinking water, n = 22
Water, Urban, Influent, n = 68
PCO2 − 13%
PCO1 − 24%

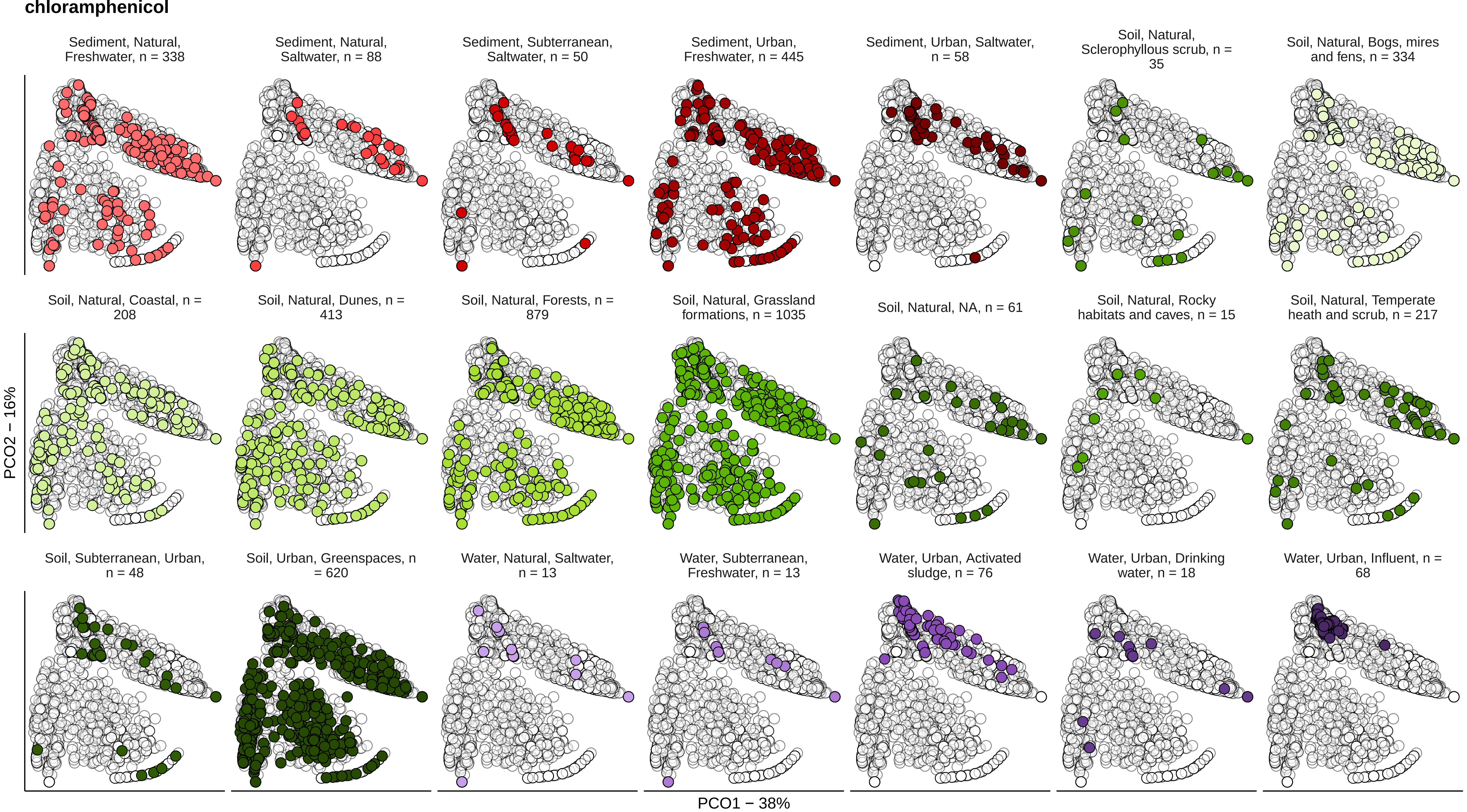

chloramphenicol
Sediment, Natural, Freshwater, n = 338
Sediment, Natural, Saltwater, n = 88
Sediment, Subterranean, Saltwater, n = 50
Sediment, Urban, Freshwater, n = 445
Sediment, Urban, Saltwater, n = 58
Soil, Natural, Sclerophyllous scrub, n = 35
Soil, Natural, Bogs, mires and fens, n = 334
Soil, Natural, Coastal, n = 208
Soil, Natural, Dunes, n = 413
Soil, Natural, Forests, n = 879
Soil, Natural, Grassland formations, n = 1035
Soil, Natural, NA, n = 61
Soil, Natural, Rocky habitats and caves, n = 15
Soil, Natural, Temperate heath and scrub, n = 217
Soil, Subterranean, Urban, n = 48
Soil, Urban, Greenspaces, n = 620
Water, Natural, Saltwater, n = 13
Water, Subterranean, Freshwater, n = 13
Water, Urban, Activated sludge, n = 76
Water, Urban, Drinking water, n = 18
Water, Urban, Influent, n = 68
PCO2 – 16%
PCO1 – 38%

**quinolone**

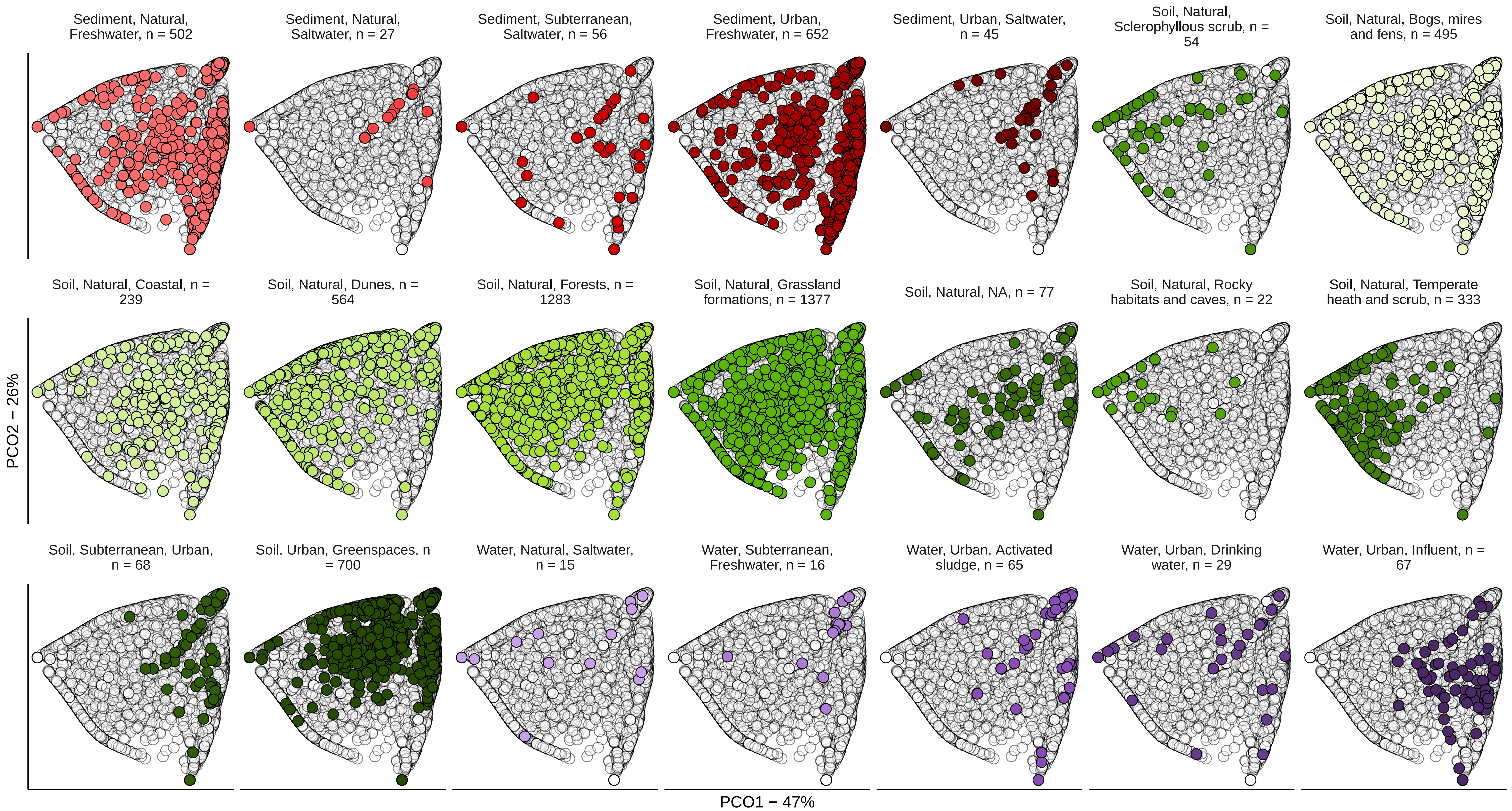

Sediment, Natural, Freshwater, n = 502
Sediment, Natural, Saltwater, n = 27
Sediment, Subterranean, Saltwater, n = 56
Sediment, Urban, Freshwater, n = 652
Sediment, Urban, Saltwater, n = 45
Soil, Natural, Sclerophyllous scrub, n = 54
Soil, Natural, Bogs, mires and fens, n = 495
Soil, Natural, Coastal, n = 239
Soil, Natural, Dunes, n = 564
Soil, Natural, Forests, n = 1283
Soil, Natural, Grassland formations, n = 1377
Soil, Natural, NA, n = 77
Soil, Natural, Rocky habitats and caves, n = 22
Soil, Natural, Temperate heath and scrub, n = 333
Soil, Subterranean, Urban, n = 68
Soil, Urban, Greenspaces, n = 700
Water, Natural, Saltwater, n = 15
Water, Subterranean, Freshwater, n = 16
Water, Urban, Activated sludge, n = 65
Water, Urban, Drinking water, n = 29
Water, Urban, Influent, n = 67
PCO2 − 26%
PCO1 − 47%

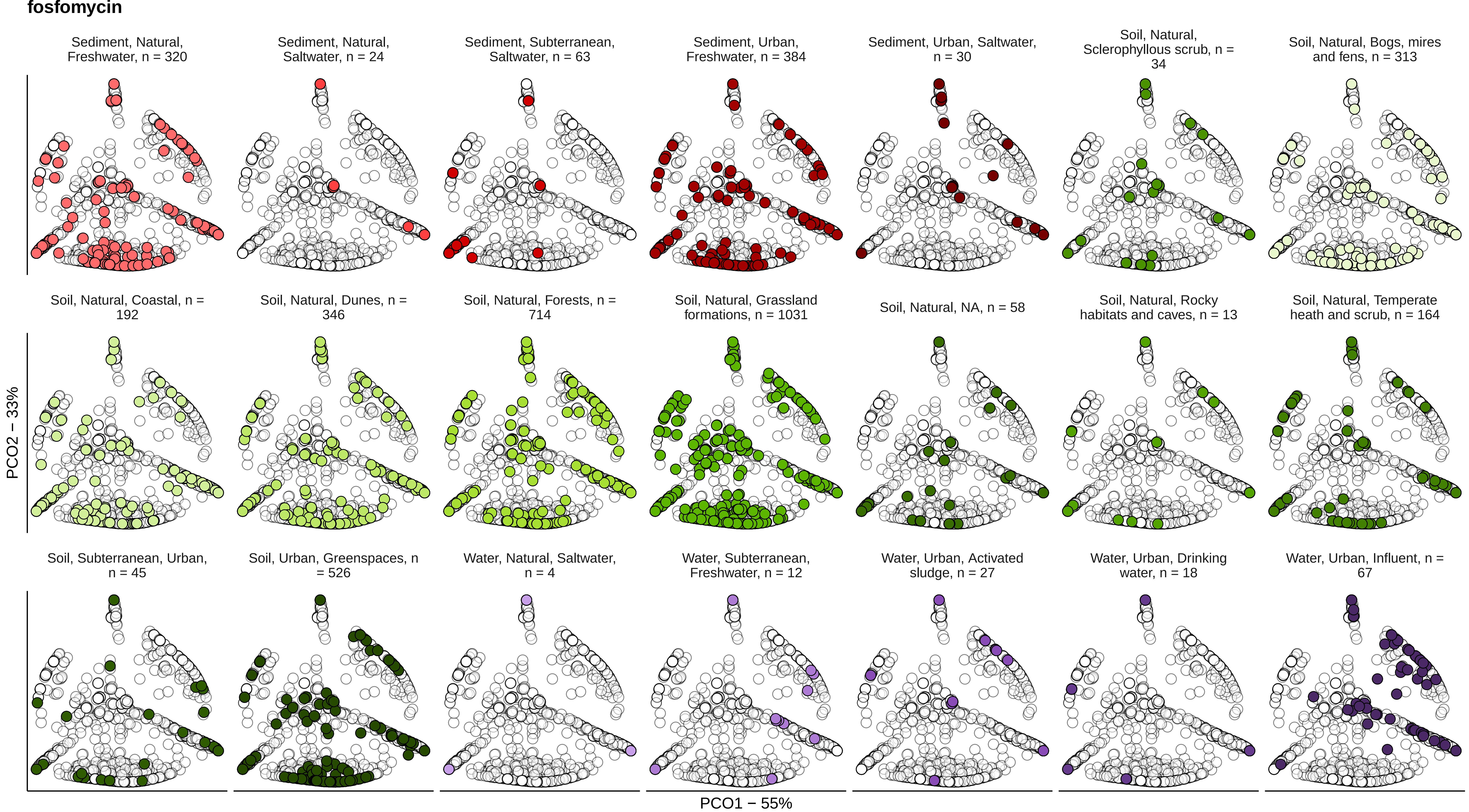
fosfomycin
Sediment, Natural, Freshwater, n = 320
Sediment, Natural, Saltwater, n = 24
Sediment, Subterranean, Saltwater, n = 63
Sediment, Urban, Freshwater, n = 384
Sediment, Urban, Saltwater, n = 30
Soil, Natural, Sclerophyllous scrub, n = 34
Soil, Natural, Bogs, mires and fens, n = 313
Soil, Natural, Coastal, n = 192
Soil, Natural, Dunes, n = 346
Soil, Natural, Forests, n = 714
Soil, Natural, Grassland formations, n = 1031
Soil, Natural, NA, n = 58
Soil, Natural, Rocky habitats and caves, n = 13
Soil, Natural, Temperate heath and scrub, n = 164
Soil, Subterranean, Urban, n = 45
Soil, Urban, Greenspaces, n = 526
Water, Natural, Saltwater, n = 4
Water, Subterranean, Freshwater, n = 12
Water, Urban, Activated sludge, n = 27
Water, Urban, Drinking water, n = 18
Water, Urban, Influent, n = 67
PCO2 – 33%
PCO1 – 55%

**bacitracin**

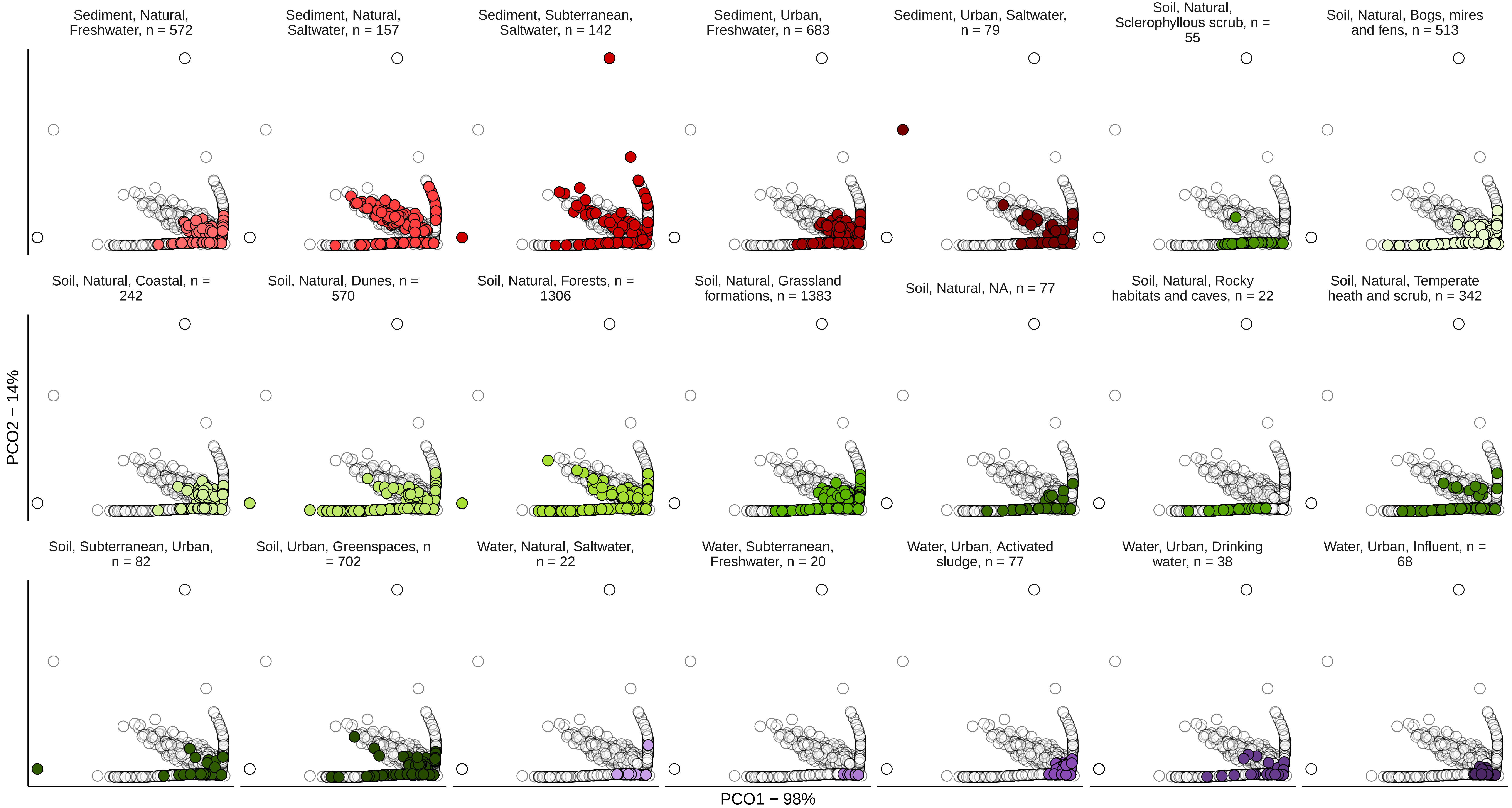
Sediment, Natural, Freshwater, n = 572
Sediment, Natural, Saltwater, n = 157
Sediment, Subterranean, Saltwater, n = 142
Sediment, Urban, Freshwater, n = 683
Sediment, Urban, Saltwater, n = 79
Soil, Natural, Sclerophyllous scrub, n = 55
Soil, Natural, Bogs, mires and fens, n = 513
Soil, Natural, Coastal, n = 242
Soil, Natural, Dunes, n = 570
Soil, Natural, Forests, n = 1306
Soil, Natural, Grassland formations, n = 1383
Soil, Natural, NA, n = 77
Soil, Natural, Rocky habitats and caves, n = 22
Soil, Natural, Temperate heath and scrub, n = 342
Soil, Subterranean, Urban, n = 82
Soil, Urban, Greenspaces, n = 702
Water, Natural, Saltwater, n = 22
Water, Subterranean, Freshwater, n = 20
Water, Urban, Activated sludge, n = 77
Water, Urban, Drinking water, n = 38
Water, Urban, Influent, n = 68
PCO2 – 14%
PCO1 – 98%

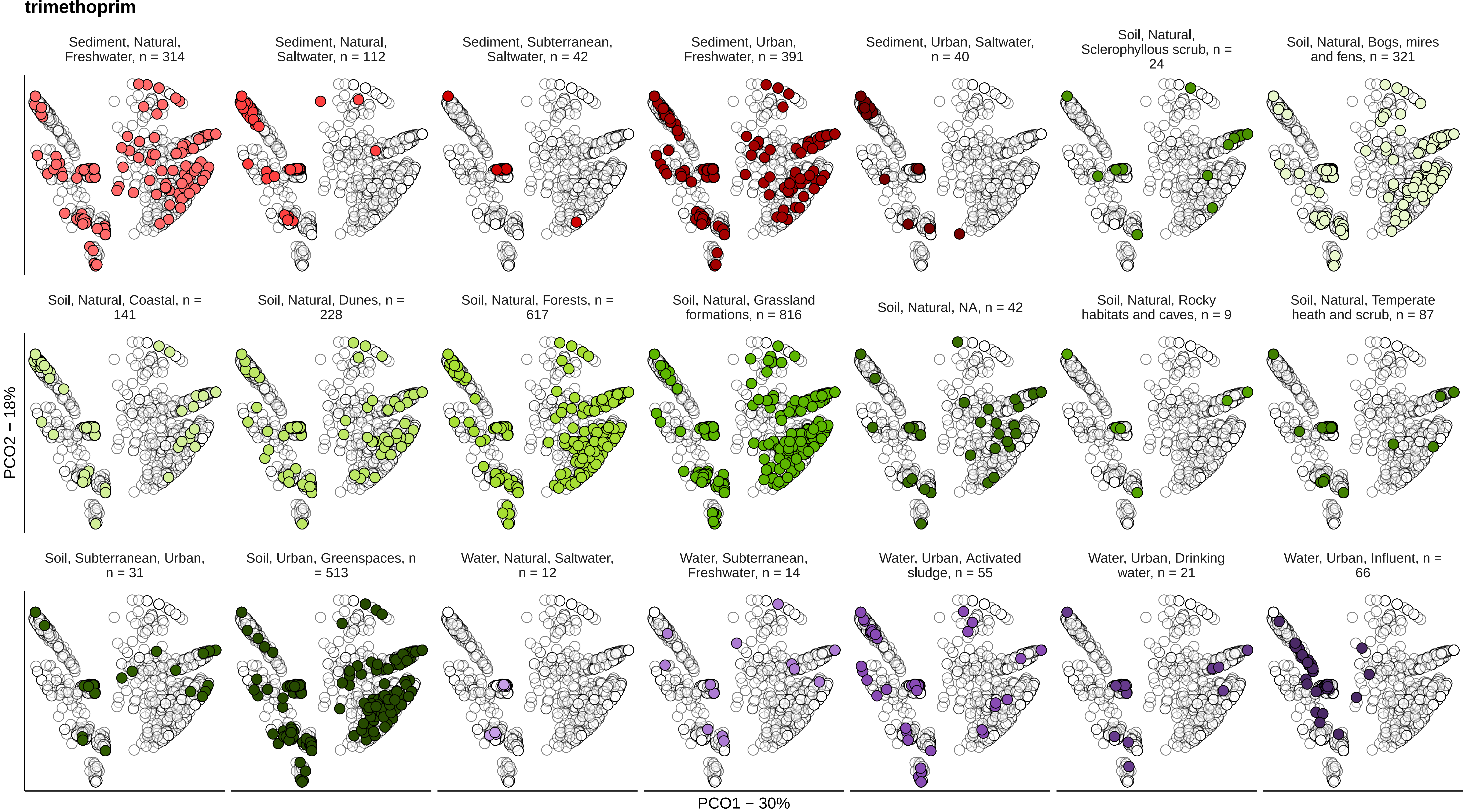
trimethoprim
Sediment, Natural, Freshwater, n = 314
Sediment, Natural, Saltwater, n = 112
Sediment, Subterranean, Saltwater, n = 42
Sediment, Urban, Freshwater, n = 391
Sediment, Urban, Saltwater, n = 40
Soil, Natural, Sclerophyllous scrub, n = 24
Soil, Natural, Bogs, mires and fens, n = 321
Soil, Natural, Coastal, n = 141
Soil, Natural, Dunes, n = 228
Soil, Natural, Forests, n = 617
Soil, Natural, Grassland formations, n = 816
Soil, Natural, NA, n = 42
Soil, Natural, Rocky habitats and caves, n = 9
Soil, Natural, Temperate heath and scrub, n = 87
Soil, Subterranean, Urban, n = 31
Soil, Urban, Greenspaces, n = 513
Water, Natural, Saltwater, n = 12
Water, Subterranean, Freshwater, n = 14
Water, Urban, Activated sludge, n = 55
Water, Urban, Drinking water, n = 21
Water, Urban, Influent, n = 66
PCO2 – 18%
PCO1 – 30%

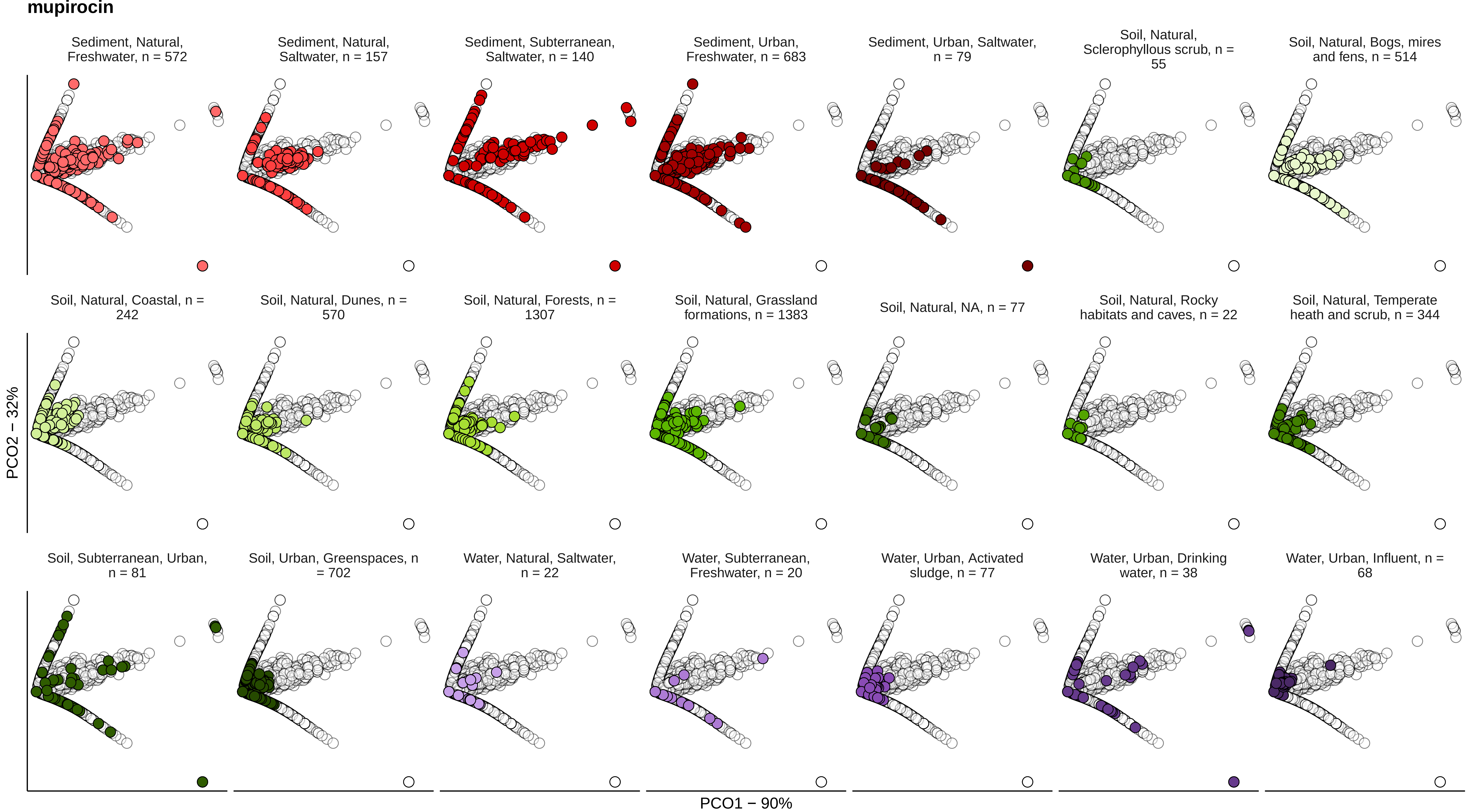
mupirocin
Sediment, Natural, Freshwater, n = 572
Sediment, Natural, Saltwater, n = 157
Sediment, Subterranean, Saltwater, n = 140
Sediment, Urban, Freshwater, n = 683
Sediment, Urban, Saltwater, n = 79
Soil, Natural, Sclerophyllous scrub, n = 55
Soil, Natural, Bogs, mires and fens, n = 514
Soil, Natural, Coastal, n = 242
Soil, Natural, Dunes, n = 570
Soil, Natural, Forests, n = 1307
Soil, Natural, Grassland formations, n = 1383
Soil, Natural, NA, n = 77
Soil, Natural, Rocky habitats and caves, n = 22
Soil, Natural, Temperate heath and scrub, n = 344
Soil, Subterranean, Urban, n = 81
Soil, Urban, Greenspaces, n = 702
Water, Natural, Saltwater, n = 22
Water, Subterranean, Freshwater, n = 20
Water, Urban, Activated sludge, n = 77
Water, Urban, Drinking water, n = 38
Water, Urban, Influent, n = 68
PCO2 − 32%
PCO1 − 90%
